# Extracellular vesicle-bound bacterial toxin pneumolysin triggers membrane engagement and damage beyond canonical pore formation

**DOI:** 10.64898/2026.03.09.710451

**Authors:** Aswathy C Sagilkumar, Avinashi Lal Kushwaha, Anuskha Shitut, Dheeraj Kumar Sarkar, Sruthika Sukumar, Jagannath Mondal, Karthik Subramanian

## Abstract

Pneumolysin (PLY) is a cholesterol-dependent pore-forming toxin and a key virulence factor of *Streptococcus pneumoniae*, the leading cause of pneumonia worldwide in children aged below 5 years. At sublytic toxin doses, host cells shed PLY-laden extracellular vesicles (EVs) during membrane repair response. However, it remains unclear how these toxin-bearing EVs engage and damage target cell membranes. Here, we combine molecular dynamics simulations, liposome fusion assays, and cell-based experiments to elucidate the membrane interaction potential of vesicle-bound PLY. Simulations indicate that EV-embedded PLY uses an exposed helix to bind target cell membranes, inducing pronounced curvature, bilayer thinning, and water influx. Liposome fusion assays demonstrate that both PLY and membrane cholesterol promote vesicle-membrane interactions. Characterization of vesicle subpopulations released from PLY-challenged monocytes by western blotting and immunogold electron microscopy demonstrated that plasma membrane-derived microvesicles are preferentially enriched in membrane-bound PLY compared to small extracellular vesicles. Consistent with these findings, MVs purified from wild-type PLY-challenged monocytes, but not the toxoid mutant PLYW433F fuse with human immune cells, delivering toxin and causing membrane damage. Together our results reveal a noncanonical, pore-independent mode of toxin dissemination, in which, vesicle-bound pneumolysin fuses and destabilize target cell membranes, representing a new proposal for EV-mediated toxin activity and a potential target for therapeutic intervention.

## Introduction

*Streptococcus pneumoniae* is a leading cause of community-acquired pneumonia and a major contributor to morbidity and mortality worldwide, particularly among children under five years of age (Infections & Antimicrobial Resistance, 2024; Kong *et al*, 2025). In 2024, the WHO classified pneumococci as one of 15 priority pathogens for which new antibiotics are urgently needed. Due to the rapidly increasing antibiotic resistance, new antimicrobial strategies targeting both the bacteria and the host need to be developed. Pneumolysin (PLY), a pore forming toxin of the cholesterol-dependent cytolysin family, is a major pneumococcal virulence determinant. It binds to cholesterol-containing host cell membranes, oligomerizes into large prepore assemblies and undergoes a dramatic conformational transition to form β-barrel transmembrane pores that drive cytolysis (Tilley *et al*, 2005; van Pee *et al*, 2017; Vogele *et al*, 2019). PLY is organized into four structurally and functionally distinct domains D1-D4 whose coordinated rearrangements drive the structural transition from soluble monomer to membrane-inserted β-barrel pore (Marshall *et al*, 2015). The membrane binding domain D4 has conserved undecapeptide residues that mediate the initial, cholesterol-dependent docking to target bilayers. Domain 3 comprises the α-helical bundles that refold into amphipathic β-hairpins that form the transmembrane β-barrel during pore insertion (Tilley *et al*., 2005). Domains 1 and 2 form the scaffold that mediates monomer-monomer contacts during prepore assembly and act as a mechanical linkage that transmits membrane-engagement to the D3 refolding event. This mechanism explains the toxin activity when PLY acts as a soluble monomer assembling directly on target cell membranes. However, increasing evidence indicates that during infection, PLY may also operate outside this classical paradigm. At sublytic concentrations, host cells activate intrinsic plasma-membrane repair pathways that sequester and remove assembled pore complexes via calcium-dependent membrane remodeling and microvesicle shedding (Wolfmeier *et al*, 2016).

The susceptibility of immune cells to PLY has been previously shown to be dependent on the microvesicle shedding capacity (Larpin *et al*, 2020). Extracellular vesicles (EVs) represent a heterogeneous population of membrane-bound particles ranging from the nano-to micron-scale particles released by cells. They include small endosome-derived exosomes (∼30–150 nm) and larger plasma membrane-derived microvesicles (∼0.1–1 μm), both of which encapsulate biological cargo such as proteins, lipids, and nucleic acids that modulate recipient cell function and mediate intercellular signalling (Jeppesen *et al*, 2023). Recently, study from our group demonstrated that PLY challenge markedly upregulates release of host EVs enriched in inflammatory cargo that mediate PLY-dependent lung injury in infected mice (Parveen *et al*, 2024). These studies establish that EV shedding is a prominent host response during pneumococcal infection and indicate that host EVs can carry bacterial toxin *in vivo.* However, the relative distribution of toxin in EV subpopulations and membrane interactions of vesicle-bound PLY was not analyzed in these studies.

Together, these observations raise the possibility that PLY presented on host-derived vesicles may represent more than a passive byproduct of cell membrane repair. Instead, vesicle-associated PLY may constitute a distinct, membrane-active form of the toxin, capable of engaging recipient membranes in ways that differ fundamentally from pore formation by soluble monomers. Because PLY oligomers embedded in a curved vesicular membrane expose a different set of protein surfaces to target cell membrane, vesicle presentation has the potential to alter toxin-lipid interactions, membrane deformation, and downstream cellular outcomes. Such a mechanism would provide a means for lateral toxin dissemination and bystander membrane damage, extending the impact of PLY beyond the primary target cell.

In this study, we test this hypothesis by combining *in silico* coarse-grained molecular dynamics simulations, *in vitro* reconstituted liposome fusion assays and functional analyses in human peripheral blood mononuclear cells exposed to purified monocyte-derived PLY-containing microvesicles. We demonstrate that PLY is more enriched in plasma membrane-derived MVs relative to small extracellular vesicles and localized on the vesicle membrane. Computational simulations indicated that vesicle-bound PLY acts as a mediator of cholesterol-dependent membrane engagement through interactions of the exposed helix with membrane lipids, inducing membrane curvature, thinning, and permeability, and facilitating toxin transfer to recipient cells. Liposome fusion, cellular fusion and membrane damage assays in human PBMCs using wild-type and mutant toxoid concur with the computational model. These findings support a revised view of pneumolysin biology in which extracellular vesicles serve as platforms for dissemination of membrane-active toxin, with important implications for pneumococcal pathogenesis and the development of host-directed therapeutic strategies.

## Results

### Computational simulation of interactions between vesicle-embedded pneumolysin and lipid bilayer

First, we performed *in silico* studies by simulating how vesicle-embedded PLY engages a target cell membrane. To capture this encounter in a controlled yet realistic setting, we built a vesicle–bilayer model system (**Figure 1**). The model comprises a coarse-grained representation (see Methods for details) of a 20 nm vesicle with an exosome-mimicking lipid composition (CHOL 40%, DPPC 30%, DOPS 20%, DPSM 10%) (Sakai-Kato *et al*, 2020). The vesicle was embedded with two PLY pentamers derived from the reported Cryo-EM structures of PLY pore complex (Vogele *et al*., 2019) (van Pee *et al*., 2017) that were oriented towards a lipid bilayer comprising of POPC 80% and CHOL 20% mimicking a target cell membrane (Rog & Pasenkiewicz-Gierula, 2006b). The PLY pentamer structure was used since it is was reported to be the minimal stable functional oligomeric form that still reflects the geometry and interactions necessary for pore initiation (Vogele *et al*., 2019). The orientation of PLY on the vesicle was fixed with the cholesterol binding D4-domain docked onto the vesicle and D1 opposed to the bilayer membrane, mimicking a vesicle containing cell-extruded PLY oligomers (Figure 1). The PLY D4 domain contains the undecapeptide loop essential for pore formation on the budding membrane. The D1 domain of PLY contains several α-helices. α-helical motifs are widely recognized for their capacity to promote membrane interactions by stabilizing protein–lipid contacts and facilitating insertion into lipid bilayers (Kabelka & Vacha, 2021). We hypothesized that the alpha helices in PLY D1 extending outward from the vesicle may play a role in membrane lipid interactions. The control system consisted of vesicle-bilayer system without any PLY oligomers (**Figure S1 and Table S1**). Both the systems were subjected to multi-microsecond long MD simulation. In the control system, no prominent changes in the bilayer or the vesicle were observed during the simulation period of 6 µs (**Figure 2A**). Biophysically, two opposing membranes are unlikely to fuse in the absence of fusion proteins due to the repulsion of the associated water molecules (Baoukina & Tieleman, 2010). However, in case of the PLY-vesicle-bilayer system, we found that pneumolysin can promote contact between the vesicle and the target membrane. Changes in membrane curvature of the bilayer was observed from 1.5 µs in the system (**Figure 2B**). This membrane curvature and its site were maintained through the simulation. By the end of 8 µs, a pronounced positive curvature was observed. To check if this was due to undulations of the bilayer or due to apposition of the vesicle, mean curvature for all three systems was conducted, suggesting that the curvature was likely induced by PLY (**Figure S2**). The curvature was observed along with membrane thinning and water permeation across the membrane (**Figures 2C-D**). This was captured in simulation snapshot shown in (**Figure 2C**) and videos (**Videos S1-S2**). Analysis of the residence time (τ) of contacts between PLY and POPC head groups calculated for last 1 µs from the 8 µs coarse-grained simulation identified amino acids in PLY domain D1 alpha helix-Leu130, Trp134 and His135 as the top interacting residues (**Figure S3)**. Residues L130, W134, and H135 were mutated to L130D, W134D, and H135G using the CHARMM-GUI mutation module to reduce local hydrophobicity and perturb the native secondary structure. (**Figure 3A** and **Figure S4**). Maximum protein-lipid contacts over the final 1 µs of simulation were analysed for both wild-type and mutant PLY. Mutation of key PLY residues led to a marked reduction in both lipid occupancy and residence time at the POPC bilayer compared with the wild-type model (**Figure 3B-C**). **Figure 3B-C** reports change in lipid occupancy relative to wild-type PLY (ΔO) and change in residence time relative to wild-type(Δτ), where larger values of ΔO and Δτ indicate a greater loss of interaction upon mutation. Notably, mutations at the most strongly interacting residues Leu130, Trp134, and His135 resulted in pronounced reduction in POPC interactions, reflected by substantially increased Δτ values and significant reductions in occupancy. These observations suggest that hydrophobic interactions at the lipid-water interface may play a critical role in stabilizing vesicle embedded PLY-membrane association. Consistently, substitution of D1 residues Leu130 to Gly130 and Trp134, His135 to Asp disrupted key membrane-anchoring interactions, leading to a marked attenuation of membrane protrusion in the mutant PLY-vesicle simulation (**Figure S5**). Additionally, no interactions and generation of curvature were observed for the mutant-PLY system.

**Figure 1.**
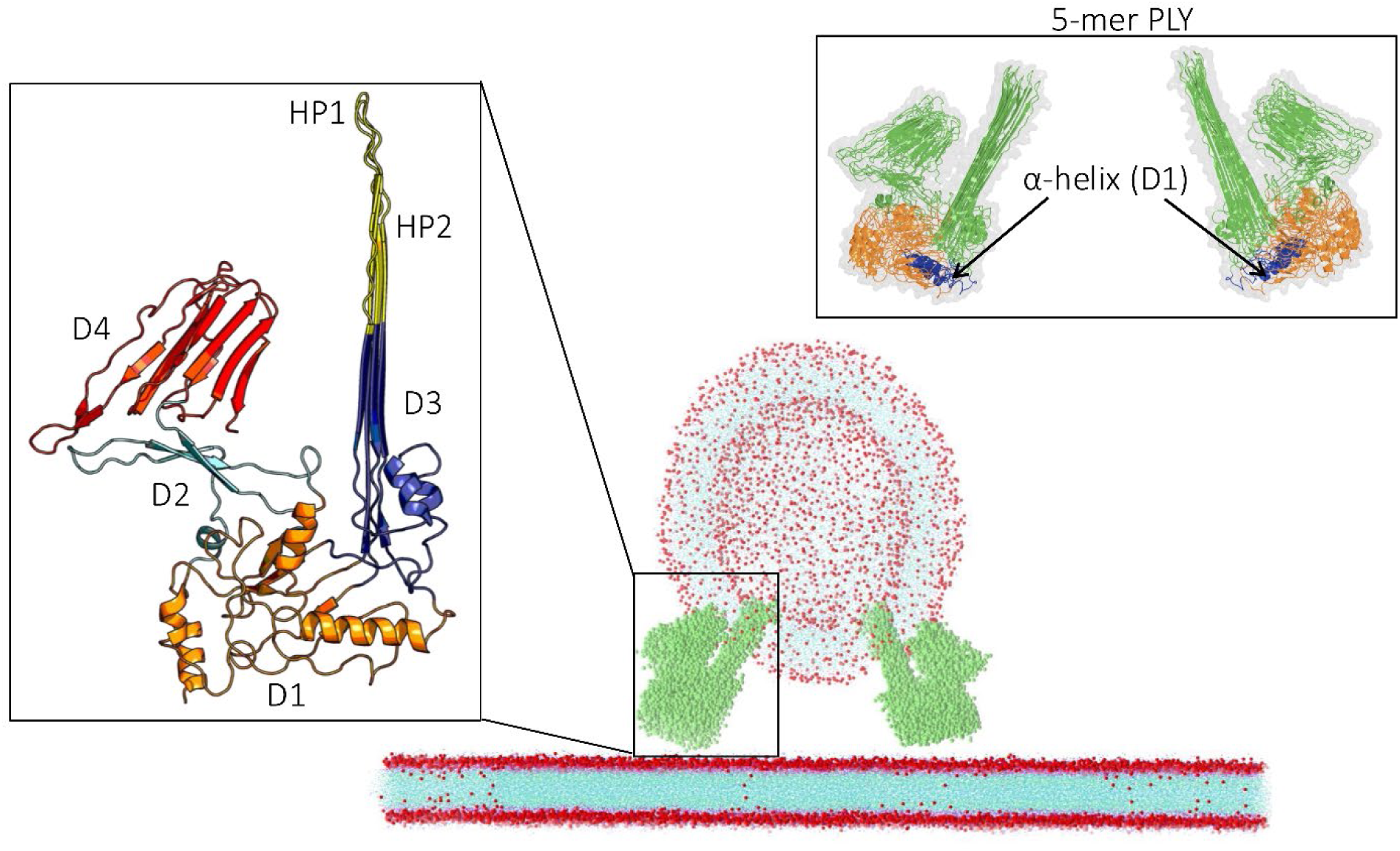
Model proposed for pneumolysin (PLY) embedded vesicle with D4-arm positioned onto the vesicle bilayer membrane and D1-arm opposed to the bilayer membrane. The 5-mer PLY structure is depicted in the cartoon and surface representation.

**Figure 2.**
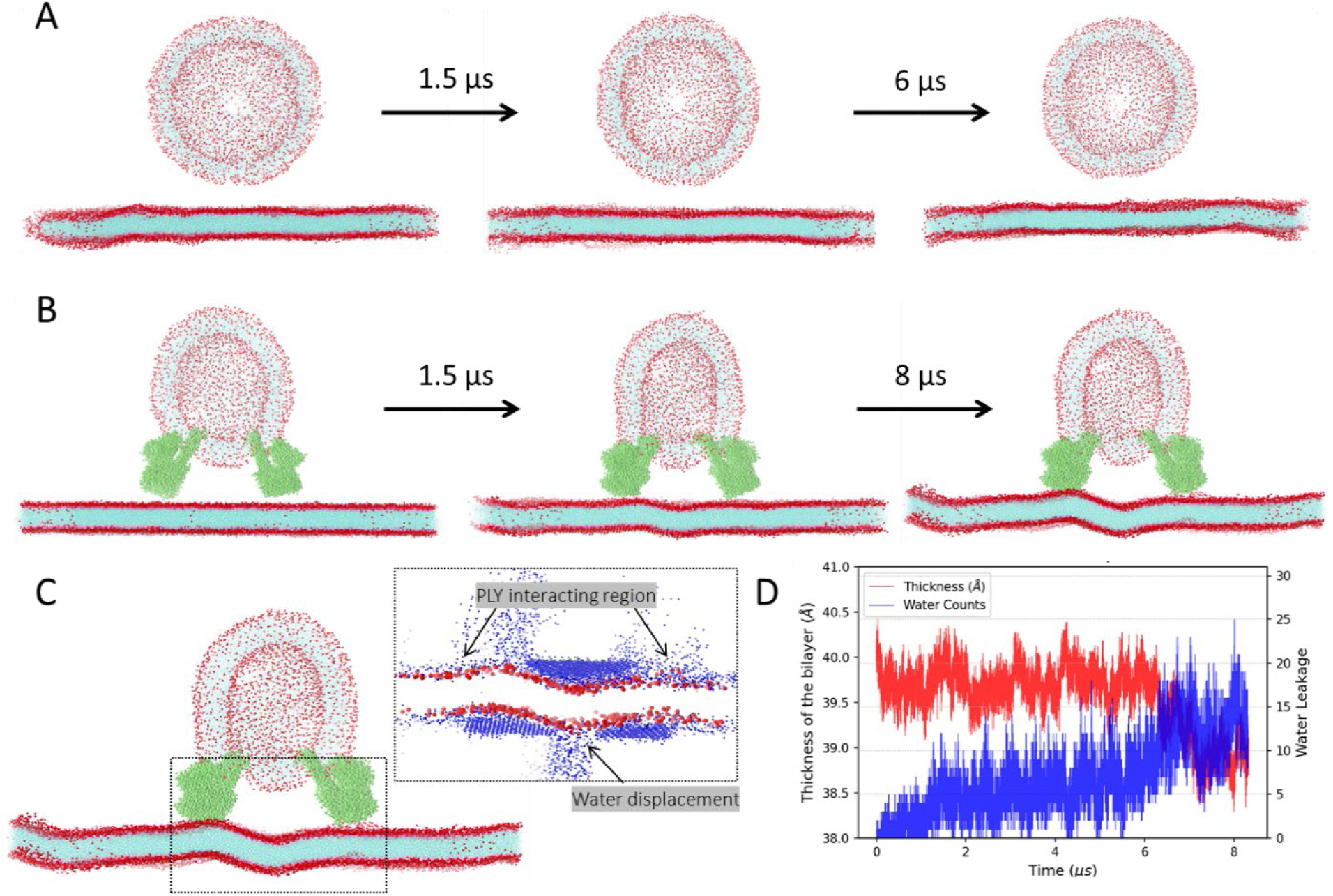
Progressive snapshots of simulation for (**A**) control with a vesicle and bilayer and (**B**) a vesicle with pneumolysin embedded and a bilayer. (**C**) Pronounced curvature was accompanied by displacement of water. (**D**) Water displacement and thickness of the bilayer were calculated for the simulation. The total time of the simulation was 6 μs for vesicle-bilayer and 6 μs for pneumolysin-vesicle-bilayer.

**Figure 3.**
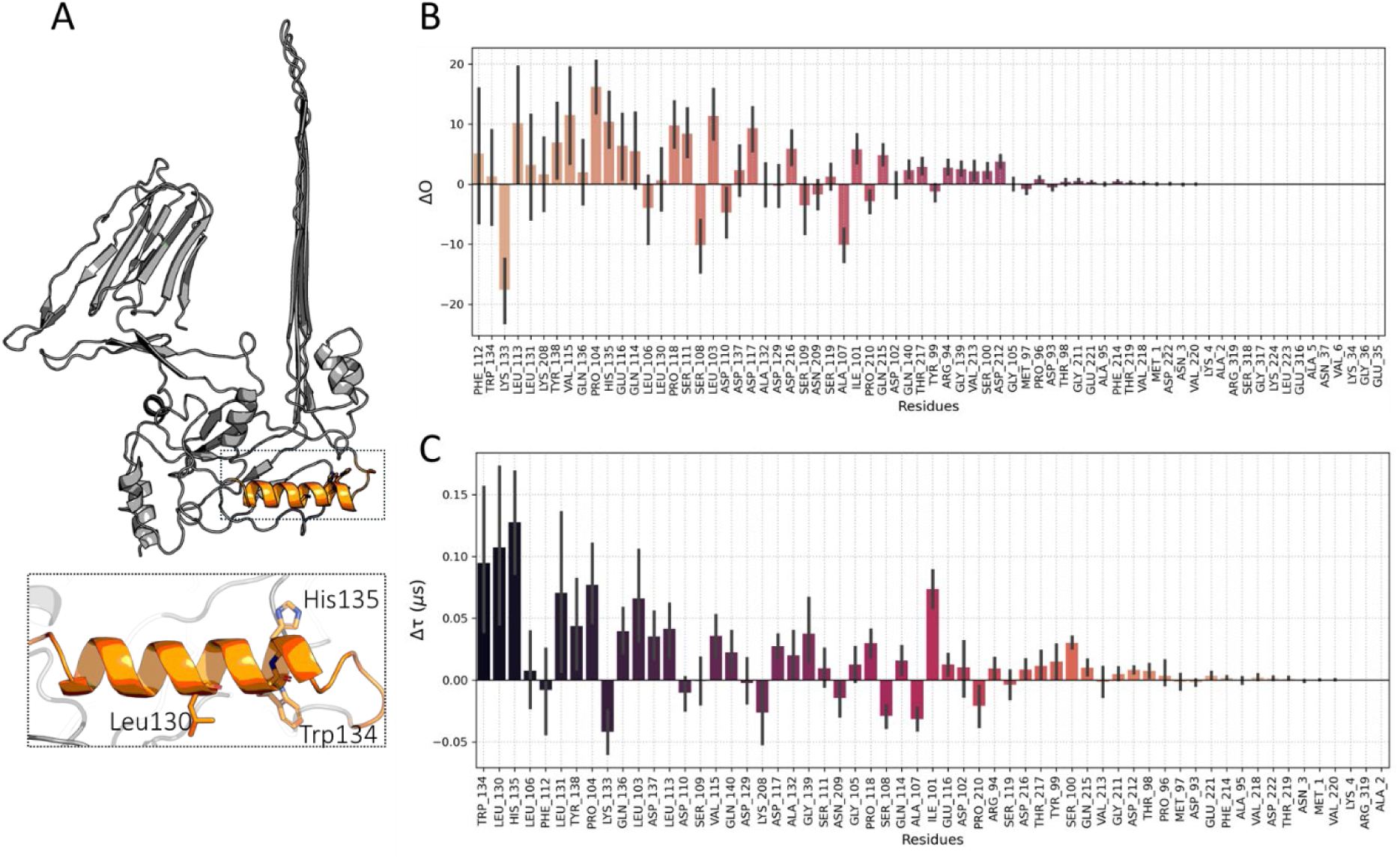
Difference in occupancy and residence time per residue of the wild type and mutant PLY. (**A**) Schematic of top interacting residues from D1 domain for which the mutations W134D, L130D and H135G were introduced. (**B**) ΔO were calculated from the difference of occupancy obtained from subtracting the mutant PLY from the wild-type PLY residues interacting with POPC, occupancy is the sum of contacts/total number of frames. The more positive the ΔO, lesser is the effect of mutant-PLY interaction with POPC. (**C**) Difference of duration (*Δ*τ) of contacts obtained by subtracting the τ of mutant PLY from wild-type PLY residue interactions with POPC. The higher the τ, the more is the duration of contact for wild-type PLY residues. The standard deviations were calculated for ΔO and Δτ plots.

Further, to study the changes in the bilayer curvature induced by PLY-vesicle, we defined a z-normal along the bilayer membrane (**Figure S6A**) and assessed the curvature of bilayer membrane for the plain bilayer, vesicle-wild-type PLY-bilayer, vesicle-mutant PLY-bilayer mutant models (**Figure S6B**). The wild-type PLY embedded vesicle-bilayer model indicated pronounced fluctuations of the bilayer, characterized by both positive and negative deviations along the membrane normal (z-axis), while the mutant PLY system showed substantially reduced bilayer perturbations, with fluctuations largely confined to positive curvature along the z-normal (**Figure S6B**). Additionally, we derived a multi-dimensional free energy surface along reaction coordinate RC-1 as bilayer thickness and RC-2 as water leakage counts. **Figure S6C** suggests a possible disruption of the bilayer induced by PLY and the progression is thermally favorable as more water leakage occurs over the simulated time. Upon quantification, the membrane thickness did not change with the mutant PLY system, however, the water leakage was similar for the wild-type PLY and mutant **(Figure S7**). Analysis of the localization tendency of this defect revealed different patterns of water displacement, prompting us to investigate how the barrier was impacted for lipid tail protrusion. Based on the unbiased simulation, it was observed that lipid tail exposure is more in the wild-type PLY system in comparison to the mutant system as inferred from the mean squared displacement and Solvent Accessible Surface Area (SASA) (**Figure S8A-B**). Taken together, the four unbiased MD simulations suggest a possible role for the Domain 1 (D1) arm of pneumolysin in modulating membrane curvature during interaction with the lipid bilayer, compared with the control and mutant systems.

### Cholesterol-dependent interactions of PLY-bound liposomes

Encouraged by the simulation’s computational prediction, we performed experiments to validate the membrane activity of vesicle-bound PLY upon interaction with recipient cells. To study the role of pneumococcal toxin, PLY during vesicle fusion, we used purified recombinant GFP-tagged PLY and synthetic 1,2-dioleoyl-sn-glycero-3-phosphocholine (DOPC) /cholesterol (70:30) liposomes prepared by thin-film hydration method. Recipient liposomes were labelled with the lipophilic dye Nile red, while the donor unlabelled liposomes were pre-incubated with GFP-PLY at 37℃ for 20 min. The non-liposome bound soluble PLY was removed by quenching with equimolar ratio of cholesterol. Confocal microscopy confirmed the binding of GFP-PLY to liposomes in a cholesterol-dependent manner, since the control cholesterol-free liposomes showed negligible GFP signal (**Figure 4A**). Subsequently, the preformed GFP-PLY liposomes were quenched with cholesterol to remove free PLY and then co-incubated with recipient Nile Red-labelled liposomes at 37℃ for 30 min. Under these conditions, multiple vesicle fusion events were observed with GFP-PLY located at the fusion interface (**Figure 4A**). To test this was due to liposome-bound PLY and rule out artifacts due to carryover of soluble non-liposome bound PLY that dissociated from preformed donor liposomes and bound to Nile red liposomes, a subset of the cholesterol quenched GFP-PLY liposomes was incubated in PBS under similar conditions for 30 min. The liposome-free supernatant was incubated with recipient Nile-red liposomes. No GFP signal was observed on the Nile-red liposomes under these conditions suggesting that the PLY transfer was primarily mediated upon liposome fusion (**Figure 4A**). Co-incubation of Nile-red liposomes with cholesterol-free GFP-PLY liposomes did not result in transfer of GFP signal, indicating the role of cholesterol in vesicle membrane fusion. In addition, the localization of GFP-PLY at the sites of contact between the donor and recipient liposomes was confirmed by 3D Z-stack confocal imaging and orthogonal projections (**Figure 4B, C**). Line intensity profile measured in the region of interest at the liposome contact sites showed the GFP signal flanked by the Nile-red signals indicating its presence at the fusion site between liposomes (**Figure 4D**). Taken together, our data suggests that the liposome bound PLY toxin can promote interaction with neighbouring vesicles upon contact.

**Figure 4.**
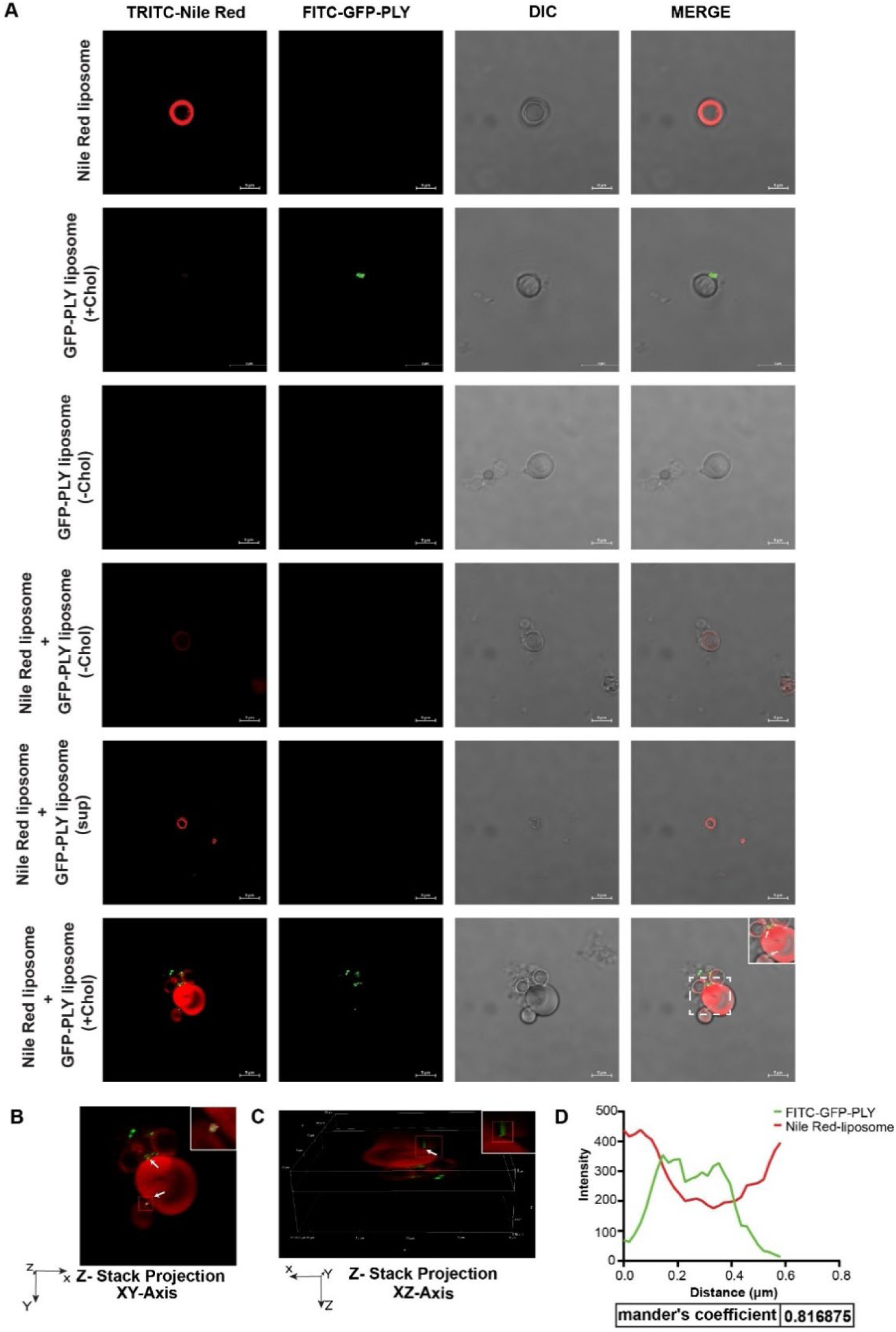
Cholesterol-dependent interactions of PLY-bound liposomes. (**A**) Confocal microscopy images of Nile Red–labeled liposomes co-incubated with GFP-PLY–bound liposomes, prepared with or without cholesterol (Chol). Scale bars, 5 µm. (**B-C**) Orthogonal projections from 3D Z-stacks in the XY and XZ axis, showing the localization of GFP-PLY at the contact sites with the recipient liposome, indicated by white arrows. (**D**). Line intensity profiles of GFP and Nile red signals were measured using Nikon A1R HD software along with the Mander’s overlap coefficient in the boxed contact area between the fusing liposomes. Data are representative of 3 independent experiments.

### Microvesicles shed from challenged human monocytes harbour PLY on the vesicle membrane

Next, we investigated the distribution of PLY in the different populations of extracellular vesicles that are shed from human THP-1 monocytes challenged with a sublytic dose of 0.5 μg/ml recombinant PLY. The microvesicle (MVs) fraction was purified by centrifugation of the conditioned culture supernatant at 10000xg. The small extracellular vesicles (sEVs) were isolated by iodixanol cushion-density gradient ultracentrifugation and pooling the fractions F5-F9. The purity of pooled sEV fractions were confirmed by probing for the tetraspanin proteins-CD63 and CD81 and absence of the major serum protein, albumin indicating purity of EVs (**Figure 5A**). The sEVs also showed the presence of PLY. Nanoparticle tracking analysis (NTA) of Naïve and rPLY sEVs revealed a mean particle diameter of 142.6 and 151.7 nm respectively (**Figure 5B**). The MVs showed expression of the classical marker for microvesicle budding from the plasma membrane marker, Annexin A1 (Jeppesen *et al*, 2019) and PLY toxin (**Figure 5C**). The MVs showed a slightly higher particle diameter compared to sEVs with mean values of 156.3 and 176.8 nm for Naïve and rPLY MVs respectively (**Figure 5D**).

**Figure 5.**
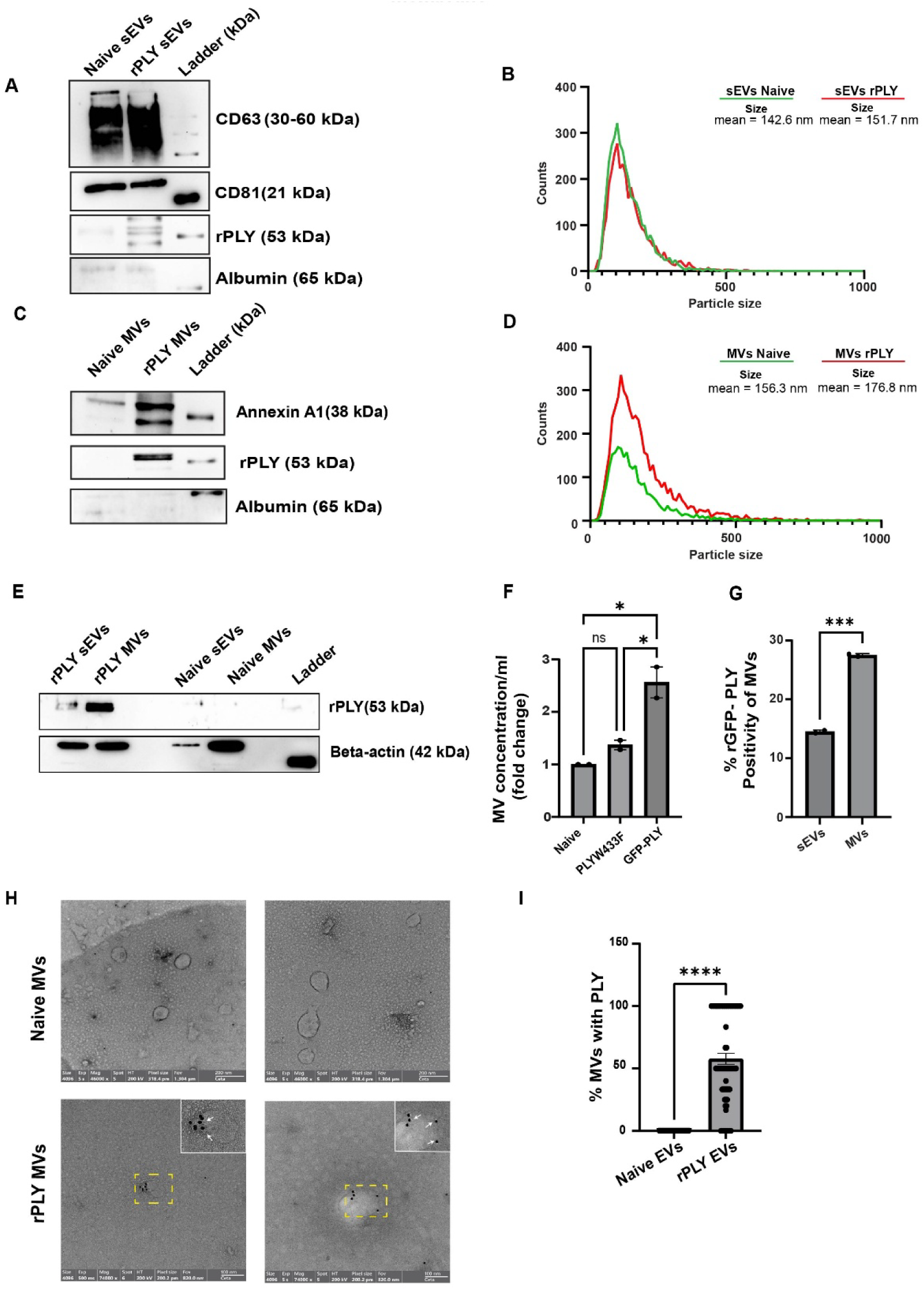
Microvesicles shed from challenged human monocytes harbour PLY on the membrane. (**A**) Western blot showing the presence of characteristic (**A**) sEV markers, CD63 and CD81 and pneumococcal toxin, PLY. Albumin was used as negative control for EV purity. **(B)** Nanoparticle tracking analysis (NTA) showing the size distribution of sEVs isolated from THP-1 monocytes challenged with (0.5 µg/ml) rPLY (rPLY sEVs) or untreated (Naïve sEVs). **(C)** Western blotting of typical microvesicle (MV) marker, Annexin A1 and PLY toxin in rPLY and Naïve MVs. (**D**) Size distribution of Naïve and rPLY-MVs using NTA. (**E**) Western blotting showing relative enrichment of PLY toxin in purified sEVs and MVs. β-actin was used as the loading control. (**F**) Quantification of MV particle concentration purified from THP-1 monocytes challenged with GFP-PLY and the mutant PLYW433F. (**G**) The percentage of MVs containing GFP-PLY was quantified using fluorescence NTA on the FL500 channel. (**H**) Representative immunogold TEM images showing the membrane localization of PLY on the MVs. Arrows indicate the (∼12 nm) gold beads conjugated to the secondary antibody, which are present on rPLY-MVs but absent on naïve MVs. Scale bars, 200 nm (Naïve) 100 nm (rPLY MVs). (**I**). Distribution of rPLY-positive MVs (N = 67 fields). The graph displays the percentage of rPLY-positive MVs and presented as the mean ± SEM. **** represents p < 0.0001 by unpaired t test. Data are representative of 3 independent experiments.

Western blot analysis revealed a markedly greater enrichment of PLY protein in the MVs compared to the sEVs, suggesting preferential incorporation of PLY protein on plasma-membrane derived MVs (**Figure 5E**). Fluorescence-mode NTA analysis of MVs isolated from GFP-PLY challenged THP-1 cells showed ∼2.5-fold increase in particle concentration compared to Naïve MVs. In contrast, MVs purified from cells challenged with the toxoid mutant, PLYW433F with reduced membrane binding and hemolytic activity (Shewell *et al*, 2014) did not show significant difference in particle concentration compared to Naïve EVs (**Figure 5F**). NTA quantification of GFP-PLY on MVs on the fluorescence mode revealed ∼2-fold enrichment of PLY in MVs relative to sEVs (**Figure 5G**) in agreement with Figure 5E. To visualize the PLY localization on the MVs, we performed immunogold labeling followed by transmission electron microscopy analysis showing membrane-bound PLY (**Figure 5H**). Quantification of TEM images revealed that ∼50% of the MVs exhibited PLY positivity (**Figure 5I**). Together, the above findings suggest that pneumococcal toxin PLY upregulates shedding of plasma membrane-derived MVs that traffic PLY toxin shed from challenged cells.

### Transfer of MV-bound PLY to recipient PBMCs upon co-incubation

We next tested whether that the PLY-bound MVs would interact and fuse with recipient cells resulting in the transfer of PLY toxin to the recipient cell membrane. Purified MVs from wild-type PLY challenged THP-1 monocytes were co-incubated with peripheral blood mononuclear cells (PBMCs) isolated from healthy donors at 37℃ for 1h. The presence of PLY in the MVs purified from challenged monocytes was validated by western blotting (**Figure 6A**). The MV-free supernatant was also probed as a control, which showed negligible carry over of free soluble PLY into EV preparations. Monocytes challenged with the PLY toxoid mutant, PLYW433F having compromised hemolytic activity relative to wild-type PLY (Paton *et al*, 1991) also shed MVs containing PLY.

**Figure 6.**
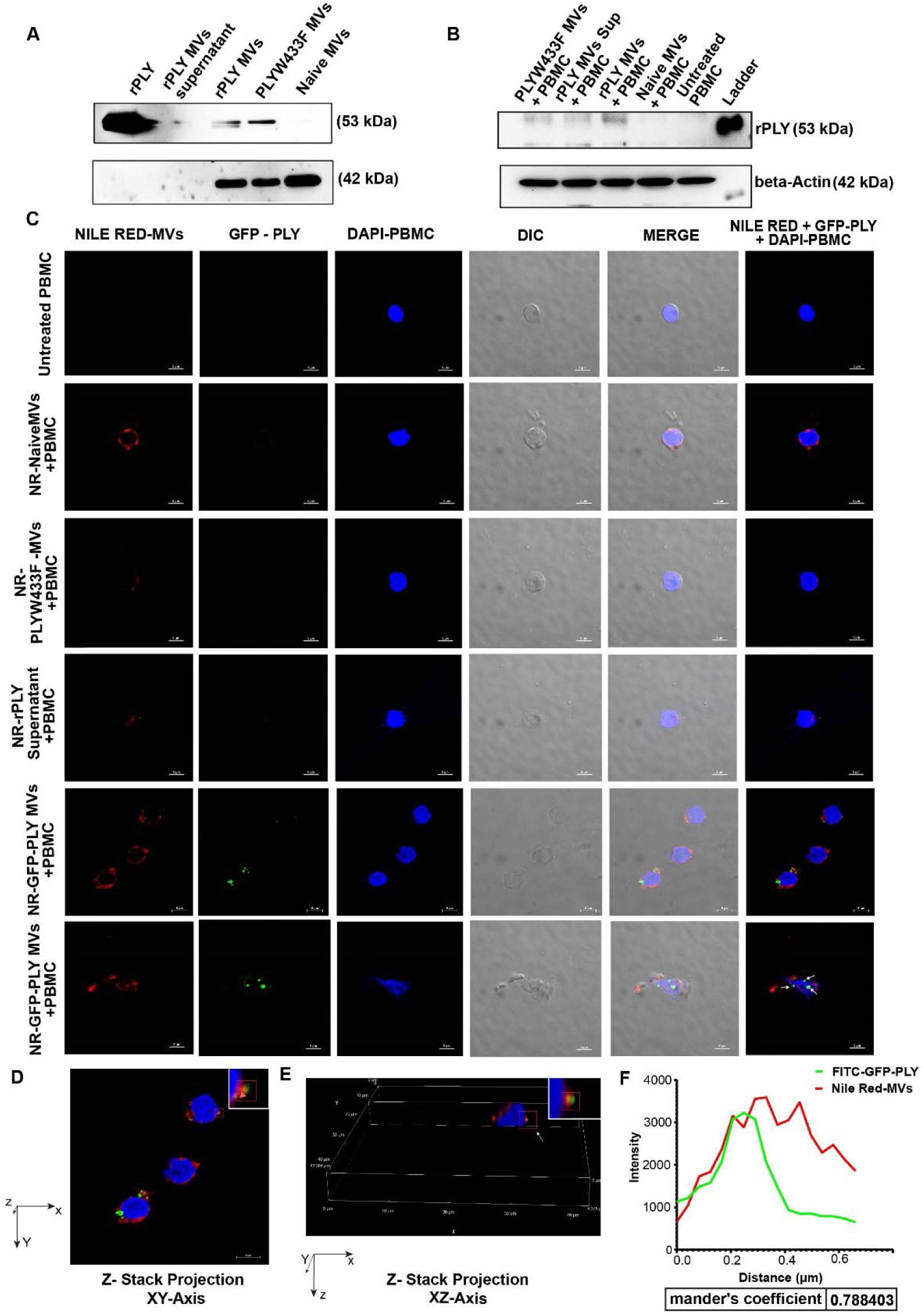
Transfer of MV-bound PLY to recipient PBMCs upon co-incubation. (**A**). Immunoblotting showing the presence of PLY in MVs purified from THP-1 monocytes challenged with wild-type PLY and the toxoid mutant PLYW433F. MV-free supernatant from PLY-MVs and naïve EVs were used as control showing negligible non-EV bound PLY after purification. β-actin served as the loading control. **(B)**. Western blot analysis of human PBMCs treated with rPLY MVs at 1h post MV incubation. PBMCs treated with wild-type PLY-MVs, but not the PLYW433F mutant or MV-free supernatant showed transfer of PLY. β-actin was used as the loading control. **(C)**. Confocal microscopy of PBMCs incubated with 30 µg of Nile Red-labelled GFP-rPLY-MVs, GFP-rPLY-MVs supernatant, PLYW433F MVs, or naïve MVs for 1h. Images show that GFP-rPLY-MVs fuse with PBMCs, showing the membrane disruption and internalization of GFP signals (arrows) on the Nile red membrane, while PLYW433F MVs and naïve MVs show no membrane damage. DAPI staining highlights the nuclei. Scale bars, 5 µm. **(D, E)**. Orthogonal projections from 3D Z-stacks in the XY and XZ plane, showing colocalization of PLY-GFP signal with MV-Nile red bound to the PBMCs. (**F**). Line intensity profiles of GFP and Nile red signals were measured using Nikon A1R HD software along with the Mander’s overlap coefficient in the selected region of interest. All data are representative of three independent experiments.

To study the transfer of MV-associated PLY to recipient PBMCs upon coincubation, we performed western blotting on the treated PBMC lysates, which confirmed the transfer of PLY on the treated cells (**Figure 6B**). Incubation with MV-free supernatant and MVs containing PLYW433F toxoid showed negligible PLY transfer to PBMCs. To visualize the MV-mediated PLY transfer on the cell membrane, we performed confocal microscopy on PBMCs incubated with MVs isolated from monocytes challenged with GFP-tagged PLY. The MVs were also counter labelled with the lipophilic dye, Nile red. Results showed that PBMCs incubated with Nile-red labelled MVs containing GFP-PLY showed co-localized Nile red and GFP signals confirming the MV-mediated PLY transfer to PBMCs (**Figure 6C**). Moreover, the cells showed ruptured cell membrane consistent with the seeding of PLY-pore. 3D Z-stack confocal imaging and orthogonal projections were also performed to confirm the membrane localization of the signals (**Figures 6D and E**). Further, line intensity profile of fluorescent signals measured at the region of interest on the PBMC membrane indicated colocalization of GFP and Nile-red signals with a Mander’s coefficient of 0.78 (**Figure 6F**). Naïve MVs isolated from untreated cells also showed binding, but the cells retained rounded and intact morphology. The MV-free supernatant as well as the MVs containing the toxoid PLYW433F showed very less cellular binding and intact cells.

### Transfer of MV-bound PLY compromises plasma membrane integrity

Next, we wanted to study the impact of MV-mediated transfer of PLY onto recipient cells. We performed propidium iodide staining (PI) assay to assess the membrane integrity of PBMCs upon MV coincubation at 37℃ for 1 h. We found that upon PLY-MV coincubation, there was a 3-fold increase in the percentage of PI positivity of treated PBMCs, relative to Naïve-MVs indicating compromised membrane integrity (**Figure 7A**). This was consistent with the higher binding of PLY-GFP with the PBMCs upon co-incubation with PLY-MV, but not Naïve MVs (**Figure 7B**). Cells treated with MVs containing the toxoid PLYW433F showed only basal level of PI positivity. The fold change in the median intensity was calculated from three independent experiments showing a clear but modestly trend (**Figure 7C**). When the plasma membrane of eukaryotic cells is mechanically injured, Ca^2+^ influx triggers a rapid repair process that promotes membrane resealing (Bhattacharya *et al*, 2022) (Andrews & Corrotte, 2018). To test whether Ca^+2^-dependent intrinsic membrane repair could reduce the impact of PLY-MVs, we next performed the experiment in the presence of 25 mM EDTA to chelate divalent cations including Ca^2+^ and Mg^2+^. We found that this caused a drastic increase in the PI positivity of treated PBMCs to 90% (**Figure 7D**) with a 4-log fold increase in the median fluorescence intensity (**Figure 7E**), suggesting the role of membrane repair process in mediating resistance to PLY-containing MVs.

**Figure 7.**
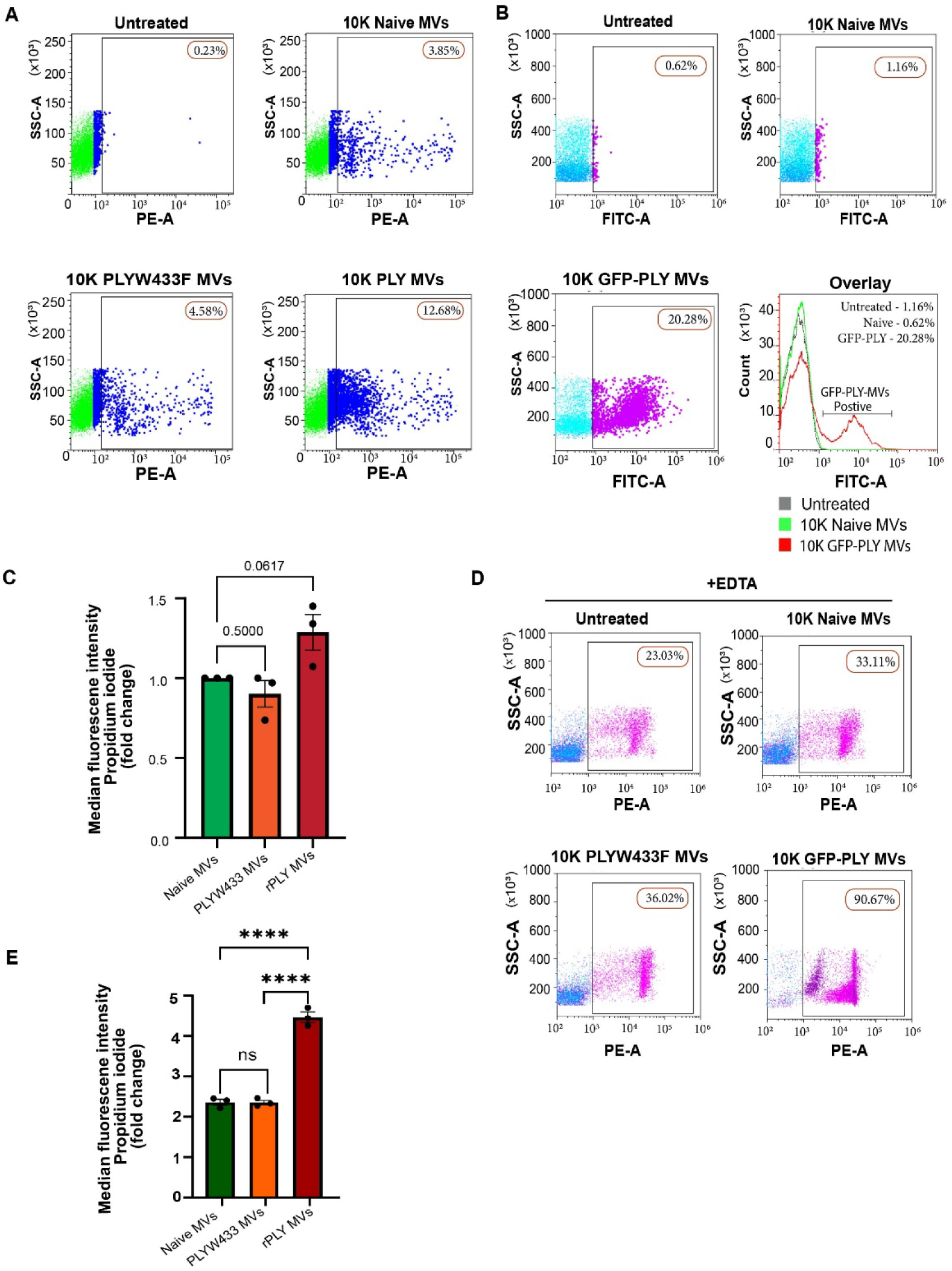
Transfer of MV-bound PLY compromises plasma membrane integrity. **(A)**. Flow cytometry histograms showing propidium iodide (PI) staining of human PBMCs at 1h post-treatment with 100 µg of rPLY MVs as measured on the PE-A channel (**B**). Flow cytometric analysis showing the percentage of GFP-PLY MVs bound to PBMC membranes after treatment with 100 µg of GFP-PLY MVs as measured on the FITC-A channel. (**C**). Quantification of median fluorescence intensity of propidium iodide uptake of human PBMCs in panel A. p-values were computed by unpaired t test. (**D**). PI staining of human PBMCs treated with 100 µg of rPLY, PLYW433F, or naïve MVs in the presence of 25 mM EDTA. (**E**) Quantification of median fluorescence intensity of propidium iodide uptake of human PBMCs treated with Naive MVs, PLYW433F, and rPLY MVs in panel D. **** represents p <0.0001 by one-way ANOVA Data are representative of three independent experiments and are presented as mean ± SEM.

Taken together, our results show that MVs shed from PLY-challenged cells implant the toxin to recipient cells upon fusion causing membrane damage.

## Discussion

This study identifies extracellular vesicle-bound pneumolysin (PLY) as an active mediator of membrane engagement and instability, extending the classical view of PLY action beyond pore formation by soluble toxin. Our findings support a previously unrecognized model in which PLY displayed on microvesicles shed from challenged host cells has unique membrane-interactive properties that may facilitate vesicle-membrane fusion and toxin transfer, thereby contributing to bystander cell damage during pneumococcal infection (**Figure 8**).

**Figure 8.**
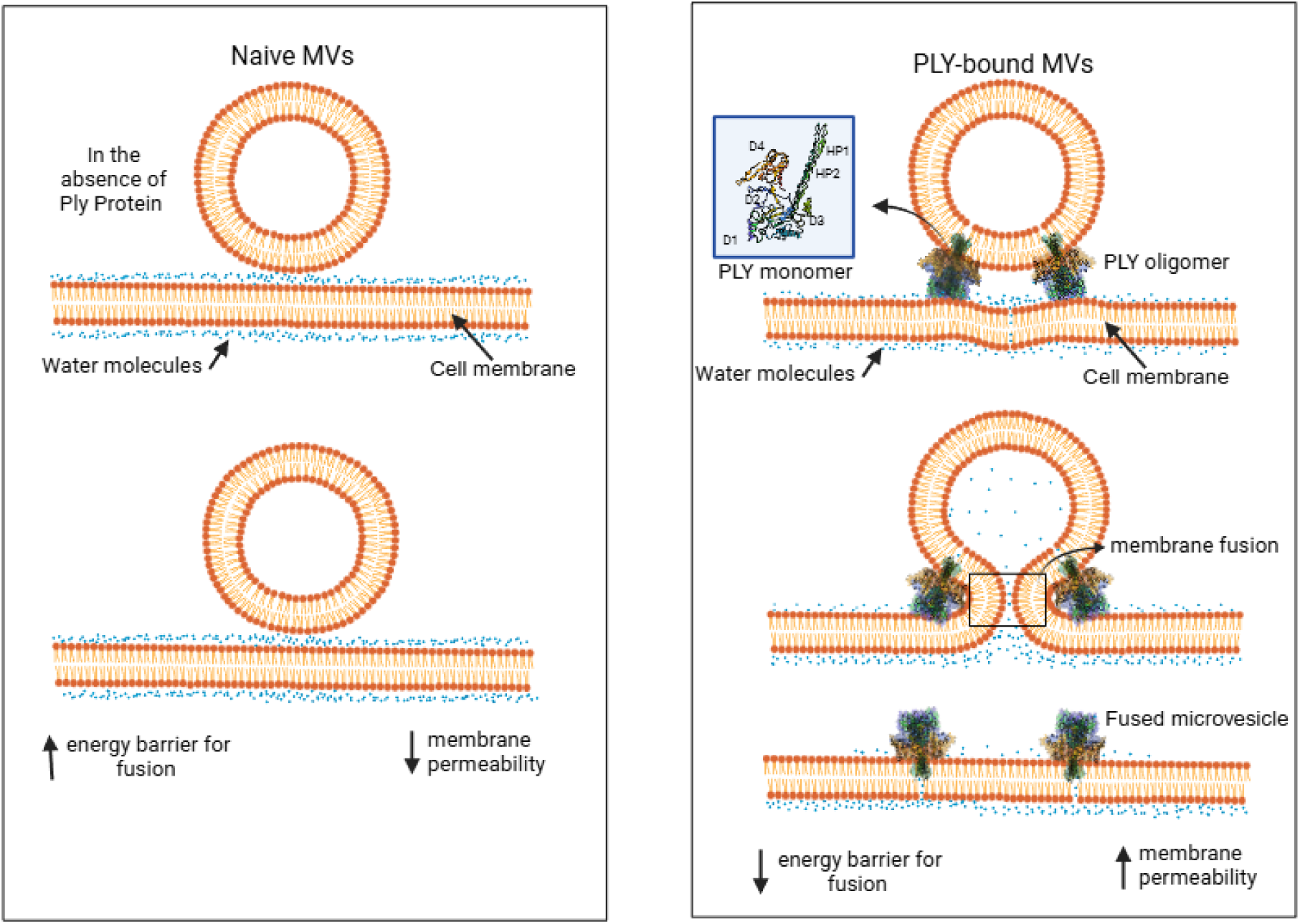
Naïve MVs show minimal fusion with opposing membranes in the absence of fusogenic proteins, likely due to a high energy barrier arising from water-mediated repulsion. In comparison, PLY-bound MVs are associated with increased membrane instability and fusion. Our work suggests that the D1 α-helix of MV-bound pneumolysin toxin may engage hydrophobically with the PBMC membrane, potentially reducing the energy barrier and facilitating membrane disruption.

Canonical models of cholesterol-dependent cytolysins (CDC) emphasize the central role of domain 4 (D4) in cholesterol recognition and of domain 3 (D3) in β-barrel pore insertion (Tilley *et al*., 2005). Our data suggest that when PLY is presented on a curved vesicular membrane, additional structural elements may become functionally relevant. In particular, the exposure of domain 1 (D1) α-helical regions enables direct interactions with an opposing bilayer, promoting membrane curvature and destabilization. This expands current understanding of CDC biology by indicating that membrane-bound oligomers can engage target membranes through non-canonical interfaces, distinct from those used during classical pore formation.

The cholesterol dependence of vesicle-vesicle interaction and toxin transfer observed here is consistent with the well-established role of cholesterol in CDC binding (Giddings *et al*, 2003) (Farrand *et al*, 2015) (Johnstone *et al*, 2022) and its ability to modulate bilayer mechanics and fusion energetics (Rog & Pasenkiewicz-Gierula, 2006a). These findings suggest that PLY may exploit cholesterol-rich membranes not only as docking platforms but potentially also as permissive environments for membrane fusion, thereby enabling lateral dissemination of toxin. At the cellular level, the preferential enrichment of PLY in plasma membrane-derived microvesicles aligns with previous work identifying microvesicle shedding as a major component of calcium-dependent membrane repair in response to CDC attack (Wolfmeier *et al*., 2016) (Larpin *et al*., 2020). While the cellular toxin shedding response is generally viewed as protective, our results indicate that the released PLY-bearing microvesicles retain pathogenic activity by transferring toxin to recipient immune cells.

This work also complements the emerging idea in the field that extracellular vesicles act as carriers of bacterial virulence factors and inflammatory mediators during infection (Jeppesen *et al*., 2023); (Parveen *et al*., 2024). Importantly, our findings provide mechanistic insight into how vesicle-associated PLY can directly engage and destabilize recipient membranes, besides acting through the reported endocytic uptake pathway (Letsiou *et al*, 2021). The reduced activity observed with the toxoid mutant, PLYW433F further underscores the requirement for intact toxin-membrane interactions even in the vesicle-bound context.

While the approaches used here provide mechanistic insight into vesicle-bound PLY-membrane interactions, they also have limitations and highlight potential scope for future work. The coarse-grained molecular dynamics simulations employed in this study are well suited to capturing mesoscopic membrane deformations and long-time scale lipid rearrangements, but higher-resolution atomistic simulations will be valuable for resolving the detailed protein-lipid interactions and transient fusion intermediates that underlie these events. Similarly, the liposome fusion assays were intentionally designed as reductionist systems to isolate the role of pneumolysin and cholesterol, extending these analyses to vesicles with more complex and asymmetric lipid compositions, and incorporating additional membrane-associated proteins, will further add knowledge on the biophysical environment of host-derived microvesicles and plasma membranes. Together, these complementary approaches integrating atomistic simulations with high-resolution structural techniques such as cryo-electron microscopy or high-speed atomic force microscopy, alongside reconstituted vesicle systems with defined lipid compositions will further advance understanding of pneumolysin-mediated vesicle-membrane interactions and their relevance in physiological and pathological contexts.

Our findings open avenues for therapeutic exploration, including targeting vesicle-membrane fusion processes or targeting membrane-active surfaces of vesicle-bound PLY as adjunctive strategies to limit pneumococcal virulence in the setting of rising antibiotic resistance.

## CRediT authorship contribution statement

**Aswathy C Sagilkumar**: Conceptualization, Methodology, Formal analysis, Investigation, Data Curation, Writing-Original Draft, Visualization. **Avinashi Lal Kushwaha**: Methodology, Formal analysis, Investigation, Data Curation, Writing - Original Draft, Visualization. **Anuskha Shitut**, **Dheeraj Kumar Sarkar**: Methodology, Data Curation, Investigation, Formal analysis, Writing **Sruthika Sukumar**: Investigation. **Jagannath Mondal**. Methodology, Formal Analysis, Writing-Review & Editing, Supervision, Funding Acquisition **Karthik Subramanian**: Conceptualization, Methodology, Writing - Original Draft, Resources, Writing - Review & Editing, Supervision, Funding acquisition.

## Acknowledgments

KS was supported by extramural funding through the INSPIRE Faculty fellowship Department of Science and Technology (DST/INSPIRE/04/2019/002238), Ramalingaswami Re-entry fellowship (BT/RLF/Re-entry/46/2020) from the Department of Biotechnology, India, and intramural funding from BRIC-Rajiv Gandhi Centre for Biotechnology, Thiruvananthapuram. JM acknowledge support from the Department of Atomic Energy, Government of India, under Project Identification No. RTI 4007. We thank Prof. Anirban Banerjee, Indian Institute of Technology, Bombay, for the kind gift of purified recombinant PLY and PLYW433F proteins. We thank Dr. Nagarjun Narayanaswamy, Scientist C, BRIC-RGCB for sharing reagents for liposome preparation. The technical assistance provided by Bioimaging facility, BRIC-RGCB is also acknowledged. Figure 8 was created with BioRender.com.

## Declaration of interests

The authors report that they have no conflicts of interest in this study.

## Materials and Methods

### Preparation of membrane systems for simulations

We constructed two membrane systems using the Martini builder in CHARMM-GUI (Jo *et al*, 2008): 1) A planar bilayer (60 × 60 nm²) of ∼4 nm thickness mimicking the lipid composition of planner bilayer membranes (POPC:80%, CHOL:20%), and 2) A liposome (∼20 nm diameter) resembling lipid vesicles (CHOL:40%, DPPC:30%, DOPS:20%, DPSM:10%) (Rog & Pasenkiewicz-Gierula, 2006b) (Sakai-Kato *et al*., 2020) were prepared. A total of four molecular systems (Table S1) were prepared viz. 1) plain bilayer membrane mimicking host cell membrane, (2) the control model with vesicle on the top of the host membrane (Figure S1), (3) vesicle embedded pneumolysin (PLY) positioned towards the host bilayer membrane at a distance of 3 nm (Figure 1) and (4) a mutant system similar to that of model 3, with PLY embedded on vesicle and oriented towards the host membrane (Figure S3). PLY pentamers (5-mer) were extracted from the Cryo-EM structure (van Pee *et al*., 2017) and docked onto the vesicle with D4 inserted into the vesicle (Figure 1) using CGSB tool (Andreasen *et al*, 2025). For preparing the molecular systems, initially the membranes were desolvated, combined using PyMOL (PyMOL Molecular Graphics System, Version 1.2r3pre, Schrödinger, LLC and resolved via the CGSB tool (Andreasen *et al*., 2025).

### Coarse grained simulations of vesicle-bilayer interactions

All coarse-grained simulations (Table S1) were performed in GROMACS 2021.4 (Abraham *et al*, 2015) with the Martini v2.2 force field (Michaud-Agrawal *et al*, 2011) for proteins, lipids, water, and ions. The ion concentration used after solvation was 0.15M NaCl. Energy minimization used the steepest descent algorithm (Berendsen *et al*, 1995) (100 kJ/mol/nm tolerance). Equilibration involved gradual restraint release, increasing time steps from 10 fs to 30 fs. Periodic boundary conditions were applied; dielectric constant was 15; Coulombic and Lennard-Jones (Marrink *et al*, 2007) cutoffs were 1.1 nm. Pressure was maintained at 1 bar using Berendsen coupling during equilibration, with isotropic coupling for production runs for all systems other than the plain bilayer where semi-isotropic coupling was used. Temperature control employed the V-rescale thermostat (Bussi *et al*, 2007). Neighbor lists were updated every 20 steps. Analysis were conducted using MDAnalysis (Michaud-Agrawal *et al*., 2011) and Prolint python modules (Sejdiu & Tieleman, 2021).

### Liposome preparation

Liposomes were prepared by dissolving 70% 1,2-dioleoyl-sn-glycero-3-phosphocholine (DOPC) with or without 30% cholesterol into 1 ml of Dichloromethane (DCM) solvent. The thin lipid film was formed using a rotary flask stirred at a 45° angle in a water bath at 40 °C. The thin film formed was further hydrated using 1 ml PBS and sonicated for 15 minutes at 37 kHz at 37 °C. The resulting liposome suspension was centrifuged at 5000×g for 5 minutes at room temperature, and the pellet was resuspended in 100 μl PBS. The liposomes were labelled with 1 mM Nile red (Invitrogen) that was added separately to the DCM solvent during the preparation stage.

### Liposome fusion assay

Liposomes with or without cholesterol were incubated with 0.5 μg (1:2000) recombinant GFP-PLY (gift from Prof. Anirban Banerji, Indian Institute of Technology Bombay) with 1 mM DTT for 20 minutes. It was centrifuged at 5000×g for 5 minutes and washed to remove unbound GFP-PLY. The pellet was topped up using free cholesterol at an equimolar concentration to PLY and incubated for 20 minutes at 37 °C to bind free soluble PLY. Following a wash, the liposomes were resuspended in PBS. Subsequently, the Nile red-labelled liposomes were co-incubated with GFP-PLY bound liposomes with or without cholesterol for 30 minutes at 37 °C. It was washed and resuspended in PBS. The GFP-PLY bound liposome with cholesterol was co-incubated with PBS for 30 minutes at 37 °C as a control to test for PLY detachment. The liposomes were then incubated with Nile red liposomes for 30 minutes at 37 °C and washed with PBS. The liposomes were analysed using the Nikon A1 confocal microscopy system on their respective FITC and TRITC channels. The acquired data were analysed by optimising on NIS AR1 HD and 3D projection, and Z-stack was performed to confirm the co-localisation of GFP-PLY on the liposomes.

### Cell culture

Human THP-1 monocytes were cultured in suspension at a density of 1×10^6^ cells/ml using RPMI 1640 medium (HiMedia) supplemented with 10% fetal bovine serum (FBS) (Invitrogen). The cells were maintained at 5% CO_₂_ and 37°C medium upon subculturing every 3 days.

### Isolation of EVs from THP-1 monocytes post PLY challenge

Briefly, 6×10^7^ THP-1 monocytes are seeded in T175 flask (Corning) with 12 ml media containing RPMI 1640 supplemented with 1% Penicillin-Streptomycin. The cells were challenged with a sublytic dose of 0.5 μg/ml of wild-type rPLY, rGFP-PLY, and mutant PLYW433F (kind gifts from Prof. Anirban Banerji, IIT Bombay) or left untreated at 37°C for 24 hours. The conditioned media collected at 24 h post-PLY challenge was subjected to differential centrifugation to remove cells (300×g, 10 minutes), cell debris and apoptotic bodies (2000×g, 20 minutes) and larger microvesicles (10,000×g, 30 minutes) at 4 °C. The resulting supernatant was filtered through a 0.45μm filter (HiMedia) to remove soluble non-EV proteins. The filtrate was layered on 2 ml of 60% iodixanol solution and subjected to cushioned density ultra centrifugation at 160,000 g for 3 h at 4 °C. The supernatant was discarded and up to 3ml of the bottom layer containing 40% iodixanol and small EVs was overlaid with 3ml of 20%, 10% and 5% sucrose-iodixanol gradient and centrifuged at 100000 g for 18h at 4 °C using SW40Ti rotor. Each 1 ml fraction was collected from the top, and F5-F9 fractions that were rich in sEVs were pooled and concentrated using 100kDa cut-off columns (Amicon). The purified EVs were stored in -80 °C until use.

### Nanoparticle tracking analysis of EVs

Nanoparticle tracking analysis (NTA) were performed using Zeta view x30 Quatt instrument equipped with four lasers, 405, 488, 520, 640 nm. The EV samples were diluted (1:1000) with 10% 1x PBS for analysis. The GFP-PLY EVs and rPLY EVs was analysed using FL500 and SC488 Channels and the particle number and size distribution were analyzed at 11 individual positions inside the measuring cell under a set sensitivity and a shutter value of 100. The data was analysed by Zetaview NTA software version 1.0.2.7.

### Western blotting

The isolated EVs were lysed using equal volume of RIPA lysis buffer (HiMedia) supplemented with Protease inhibitor cocktail (Roche) on ice for 30 minutes, followed by centrifugation at 16,000 x g for 30 minutes at 4 °C. Total EV lysate protein was quantified using Bicinchoninic acid assay (Pierce). 30 μg of total EV protein was denatured using 1x Laemmli buffer (Biorad) containing 50 mM DTT and boiled at 70°C for 10 minutes and separated using 12-15% SDS polyacrylamide gel electrophoresis and transferred onto PVDF membrane. The membrane was then blocked with 5% BSA at room temperature for 1 h and and probed with mouse anti-CD81 (1:1000), rabbit anti-CD63 (1:1000), rabbit anti-Annexin A1 (1:1000) and mouse anti-PLY (1:500) incubated overnight at 4°C, followed by HRP-conjugated secondary antibody HRP-conjugated anti-mouse IgG and anti-rabbit IgG (1:3000, Biorad). The chemiluminescence was visualised with an iBright FL1500 Imaging System (Invitrogen). Subsequently, the membrane was stripped and reprobed for the loading control, using a mouse anti-β-actin antibody at a dilution of 1:1000 as the primary antibody, followed by a goat anti-mouse antibody at a dilution of 1:3000 as the secondary antibody. β-actin served as an internal control to confirm uniform protein loading across the samples.

### Isolation of PBMCs from healthy donors

20 ml of venous blood was drawn from healthy volunteers in EDTA vacutainer tubes (BD Bioscience) to prevent clotting. Informed consent was obtained in accordance with the ethical committee approved protocol (Ref. IHEC/12/2020_E/14). The collected blood was then diluted with equal volume of recommended media (endotoxin-free PBS containing 2% FBS and 1mM EDTA. The diluted blood was carefully layered over HiSep density gradient media (HiMedia) and centrifuged at 1200×g for 20 min brakes off. The buffy coat layer was carefully collected to a new tube and topped with recommended media followed by centrifugation at 300xg for 10 min with low brakes. The washes were repeated thrice. A low spin at 120xg for 10 minutes with low brakes was performed in between the washes to remove platelets. Residual RBCs were lysed with 1x RBC lysis buffer (HiMedia) for 10 minutes at room temperature. The PBMC pellet was resuspended in RPMI containing 2.5% EV-depleted FBS (Gibco), counted and seeded in 12 well plate for EV challenge.

### Propidium Iodide staining

Propidium Iodide (PI) was used to measure the membrane permeability induced by PLY-bound MVs on PBMCs. 100 µg of MVs, including GFP-rPLY, GFP-rPLY supernatant, PLYW433F, and naive EVs were co-incubated with 2 × 10^5^ PBMCs for one hour at 37°C. After incubation, unbound EVs were removed by wash and centrifugation for 5 minutes at 2,000 × g. The cell pellet was resuspended in 200 µL of FACS buffer and subsequently transferred to a 96-well plate. The PBMCs were incubated with 10 µM PI stain (Invitrogen) for 30 minutes at 37°C. Samples were washed thrice by flicking the supernatant at 2,000 × g for 1 minute. The pellet was resuspended in 200 µl of FACS buffer and analyzed on the PE-A channel on the BD FACS Aria II flow cytometer (BD Biosciences) to quantify the percentage of PI positive population. 5% H_2_O_2_ was used as a positive control to induce membrane damage to PBMCs. The data obtained were further analyzed using Kaluza software (Beckman Coulter).

### Immunogold labelling of MVs with PLY antibody

5 μl of the EV rich fractions were used for immunogold labeling using transmission electron microscopy. The samples were fixed with 2% paraformaldehyde (Himedia) to preserve the EV membrane integrity and10 μl of fixed EV samples was adsorbed onto formvar coated 400-mesh copper grids (Electron Microscopy Sciences) overnight at 4 C^ͦ^ in a humidified chamber. The excess sample was removed using a wick and blocked with filtered 0.1% BSA in PBS for 30 minutes at room temperature. The grid was incubated with 10 μl of primary antibody against PLY (1:10) for 1 hour at room temperature. The grids were washed thrice with PBS and incubated with 10 μl of 12 nm gold conjugated anti-mouse antibody (1:10) for 30 minutes at room temperature. The samples were fixed using 2.5% glutaraldehyde for 10 s, washed twice with de-ionized water and negatively stained using UranyLess (Electron Microscopy Sciences) for 10 s. The excess stain was removed using a wick and allowed to dry. The grids were then examined at 200kV and 74k magnification using Talos F200i transmission electron microscope (Thermo Scientific).

### PLY-MV fusion with PBMCs

30 µg of Naïve, PLYW4333F, GFP-PLYMVs supernatant, and GFP-PLY MVs were fluorescently labelled with Nile-red (10 µg/mL) and incubated for 30 minutes at room temperature. The samples were centrifuged at 10,000 × g for 30 minutes to remove excess dye, followed by two consecutive washes with PBS. The labelled MVs were co-incubated with 7.5 × 10^5^ PBMCs for 1 hour at 37°C. Following the incubation, the cells were washed at 300 × g for 10 minutes to remove unbound MVs. The final cell pellet was resuspended into 30 µL of PBS, and glass slides were mounted using ProLong Antifade Glass Mounting media (Invitrogen) with DAPI. Confocal imaging was performed using a laser-scanning confocal microscope, and the images were processed and analyzed using Nikon AR1HD software to assess fluorescence signals and observe the localization of the GFP-PLY protein on the PBMC membrane.

### Statistical analysis

Data were statistically analyzed using GraphPad Prism v.9.1.2. Data represent mean ± SEM unless otherwise specified. Experiments were performed with three biological replicates with triplicates unless specified otherwise. Pairwise comparison of normalized data was analyzed using unpaired t-tests. Differences were considered significant at ∗*p* < 0.05., ∗∗*p* < 0.01; ∗∗∗*p* < 0.001, ∗∗∗∗*p* < 0.0001.

## Notes

### Competing Interest Statement

The authors have declared no competing interest.

