## Supplementary figures and legends for "Extracellular vesicle-bound bacterial toxin pneumolysin triggers membrane engagement and damage beyond canonical pore formation"

**Table S1**. Complete list of all the production simulations performed

| SL no. | Simulation time (us) | Number of particles | Lipid composition | Protein | Model system |
|---|---|---|---|---|---|
| **1.** | 8 | 1588090 | POPC, CHOL, DPPC, DPSM,DOPS | yes | Ves-BL-PLY |
| **2.** | 8 | 186658 | POPC, CHOL, DPPC, DPSM,DOPS | no | Ves-BL |
| **3.** | 8 | 62683 | POPC, CHOL | no | BL |
| **4.** | 8 | 1588207 | POPC, CHOL, DPPC,DPSM,DOPS | yes | Ves-BL-mPLY |

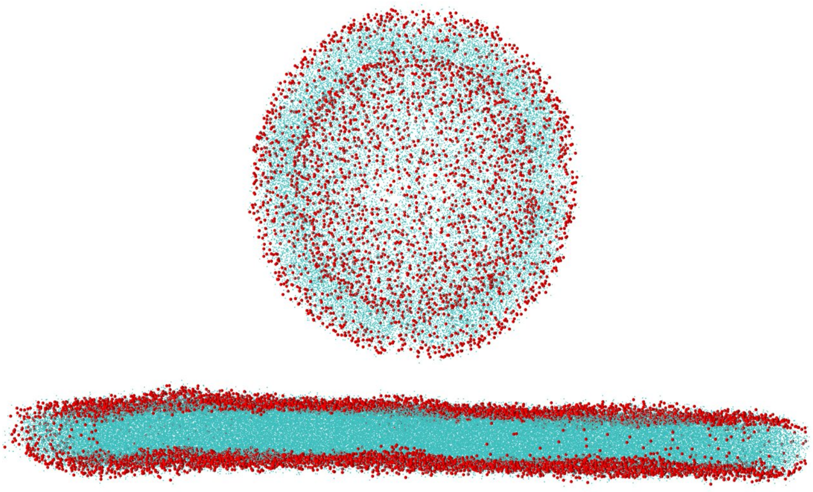

**Figure S1**. Model for control system containing a bilayer kept at 3 nm distance from the vesicle.

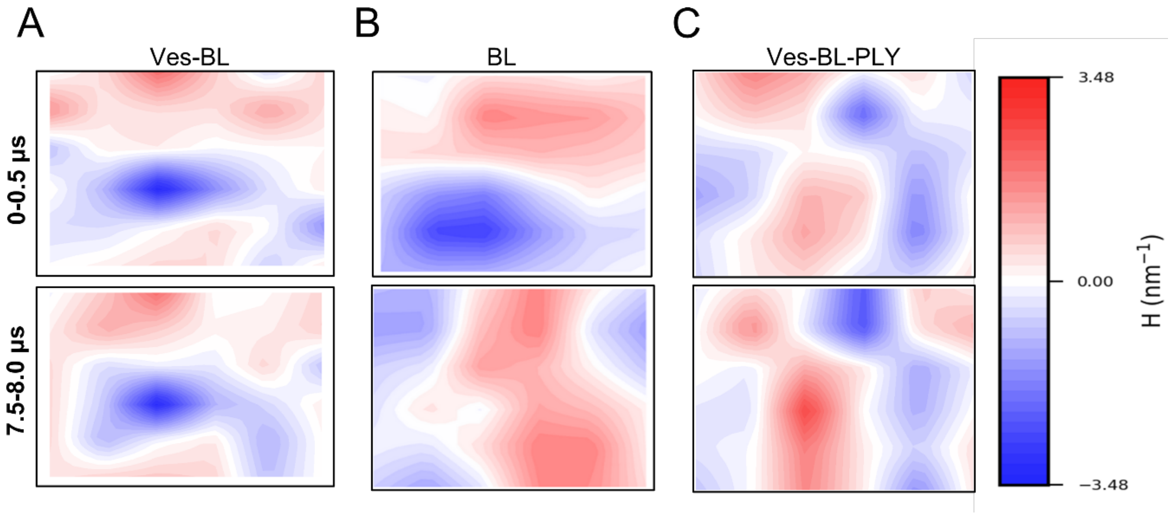


**Figure S2**. Mean curvature plots for A) Control system with vesicle and bilayer B) Plain bilayer C) Pneumolysin embedded vesicle and bilayer. This was done for 0 to 0.5 µs and 7.5 to 8 µs for the three systems.

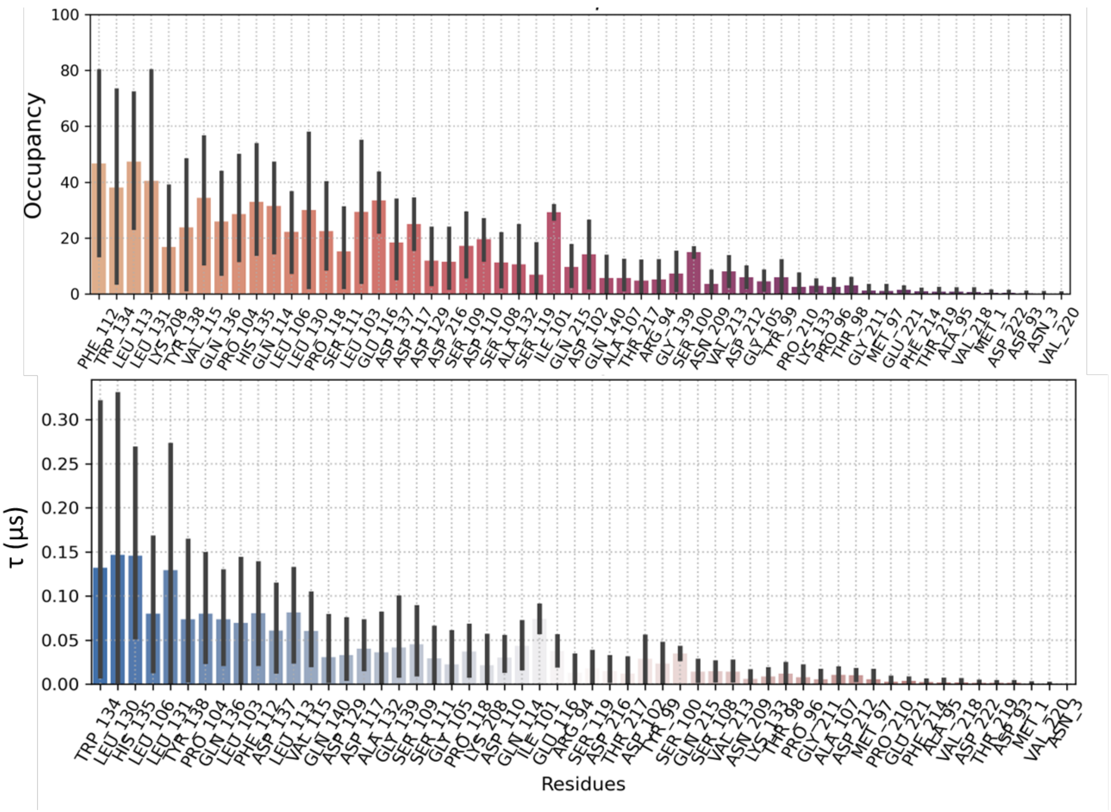


**Figure S3**. Occupancy and residence time (τ) of contacts between PLY and POPC head groups calculated for last 1 µs from 8 µs coarse-grained simulation.

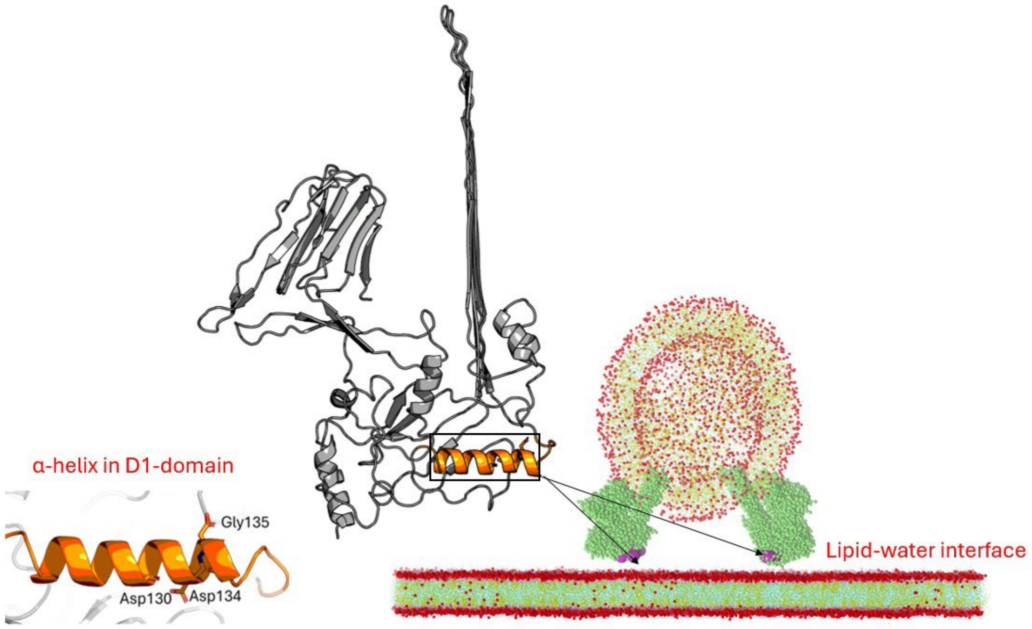


**Figure S4**. Model with pnemolysin mutations W134D, L130D and H135G with the structure of the target domain depicted.

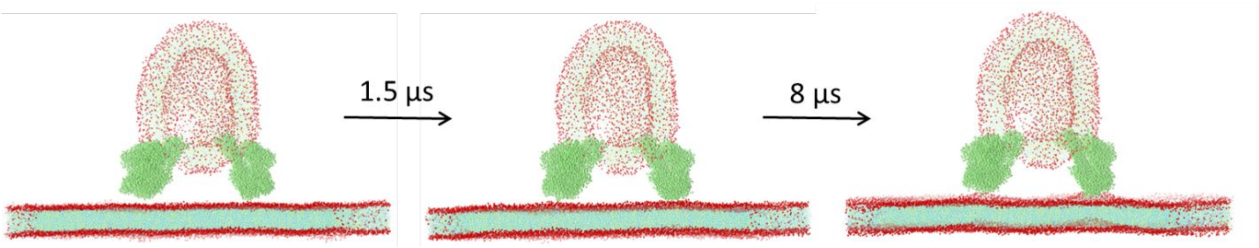


**Figure S5**. Progression of mutant PLY (CG beads shown in lime) interaction with host bilayer membrane. Snapshots from 0 μs, 1.5 μs and 8 μs are represented. The phosphate head groups are shown in red spheres for the bilayer and vesicle.

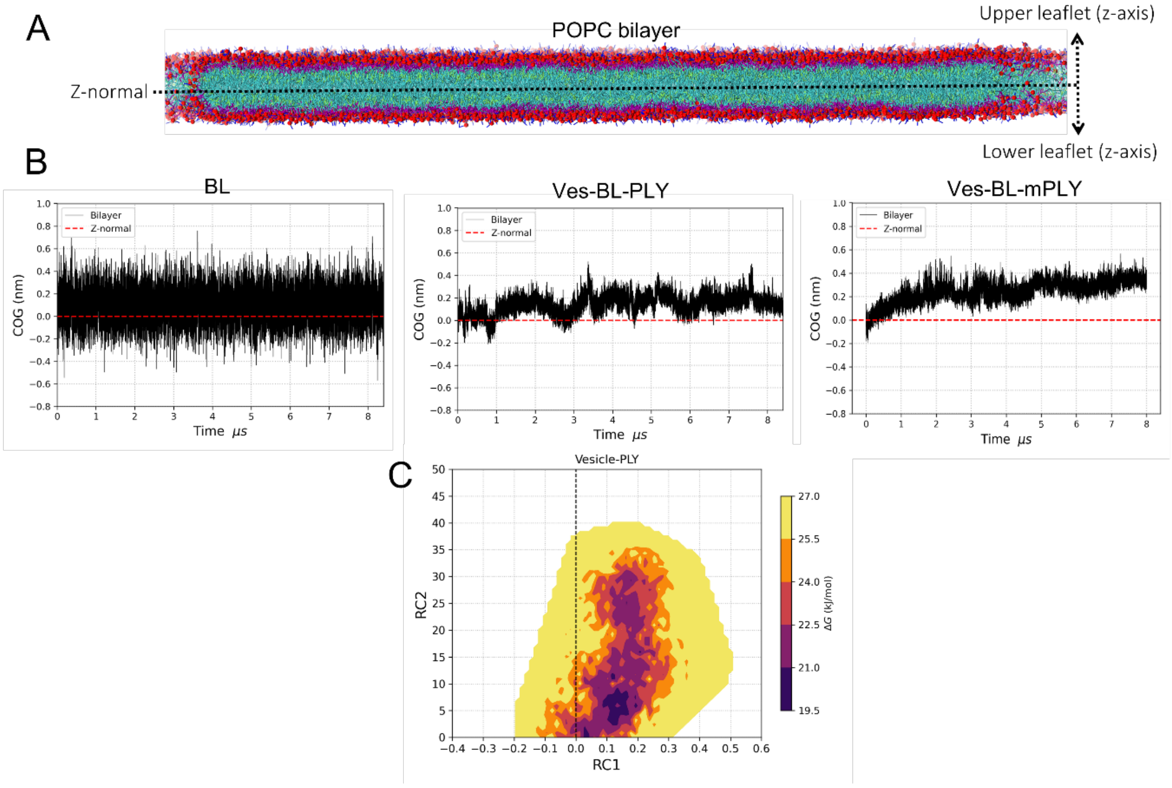


**Figure S6**. (A) Center of geometry for the membrane normalized to the center of the bilayer in X axis. (B) Center of geometry (COG) calculations with respect to z-normal (red dashed line) were done for Plain Bilayer, Bilayer with pneumolysin embedded vesicle and Bilayer with mutant pneumolysin embedded vesicle. (C) 2D Free Energy Surface to calculate the free energy between RC1 depicting the shift in center of geometry and RC2, which is the water leakage for the system with pneumolysin embedded vesicle and bilayer.

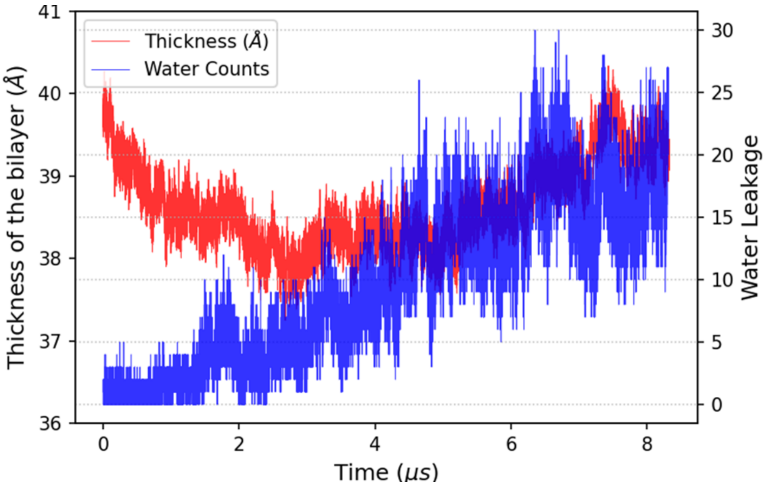


**Figure S7**. Water leakage and bilayer membrane thickness profile as a function of time for mutant PLY embedded vesicle and bilayer model.

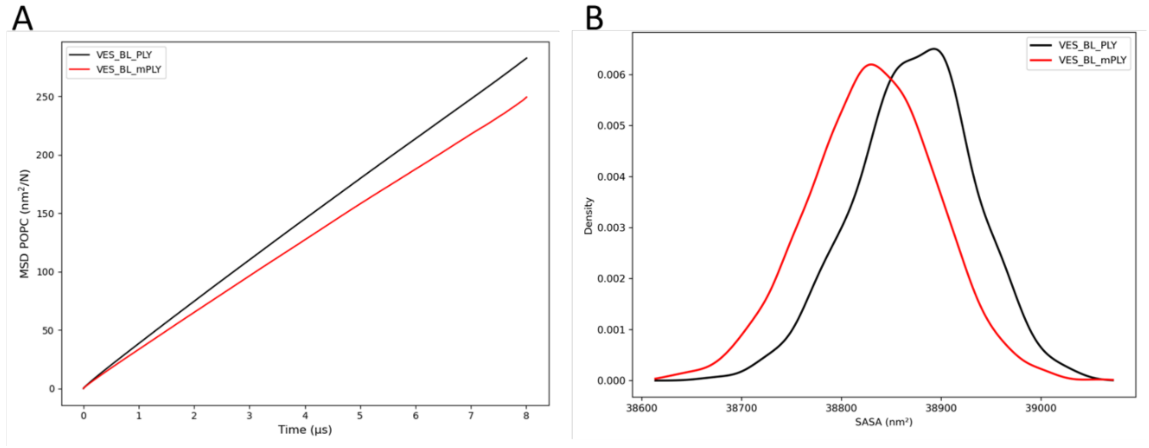


**Figure S8**. (A) Mean square displacement (MSD) and (B) Solvent Accessible Surface Area (SASA) for wild-type PLY embedded vesicle and bilayer model (in black) and mutant PLY embedded vesicle and bilayer model (in red). MSD reflects lipid mobility, with higher values indicating faster diffusion. SASA measures tail exposure: increased tail SASA corresponds to transient hydrophobic exposure and lipid packing defects that promote stalk formation, while lower SASA reflects tighter packing and a more stable bilayer. The calculations were performed for 8 µs of trajectory data with a stride of 10 frames.
