## Supplementary figures and images for "Extracellular vesicle-bound bacterial toxin pneumolysin triggers membrane engagement and damage beyond canonical pore formation"

### Figure S1

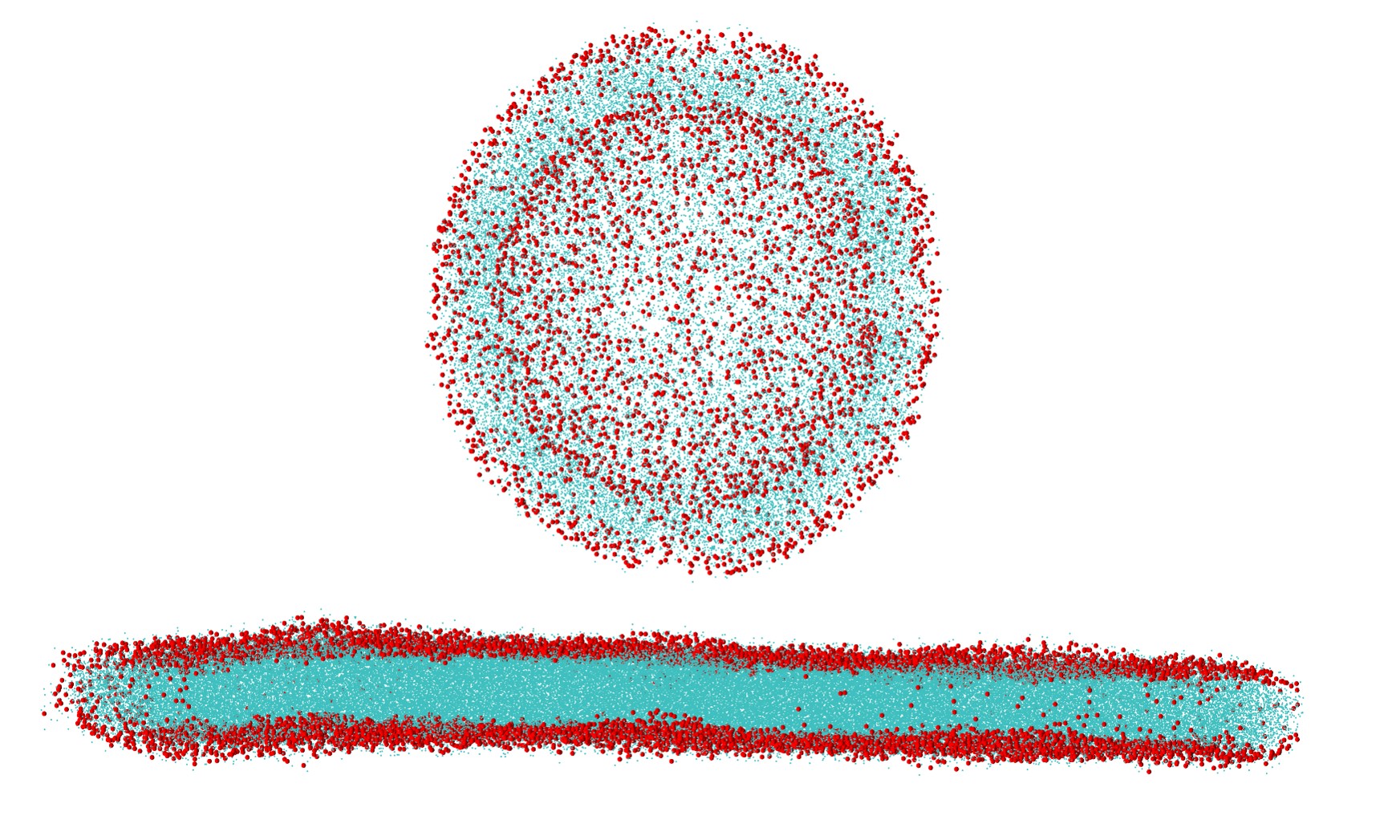

### Figure S2

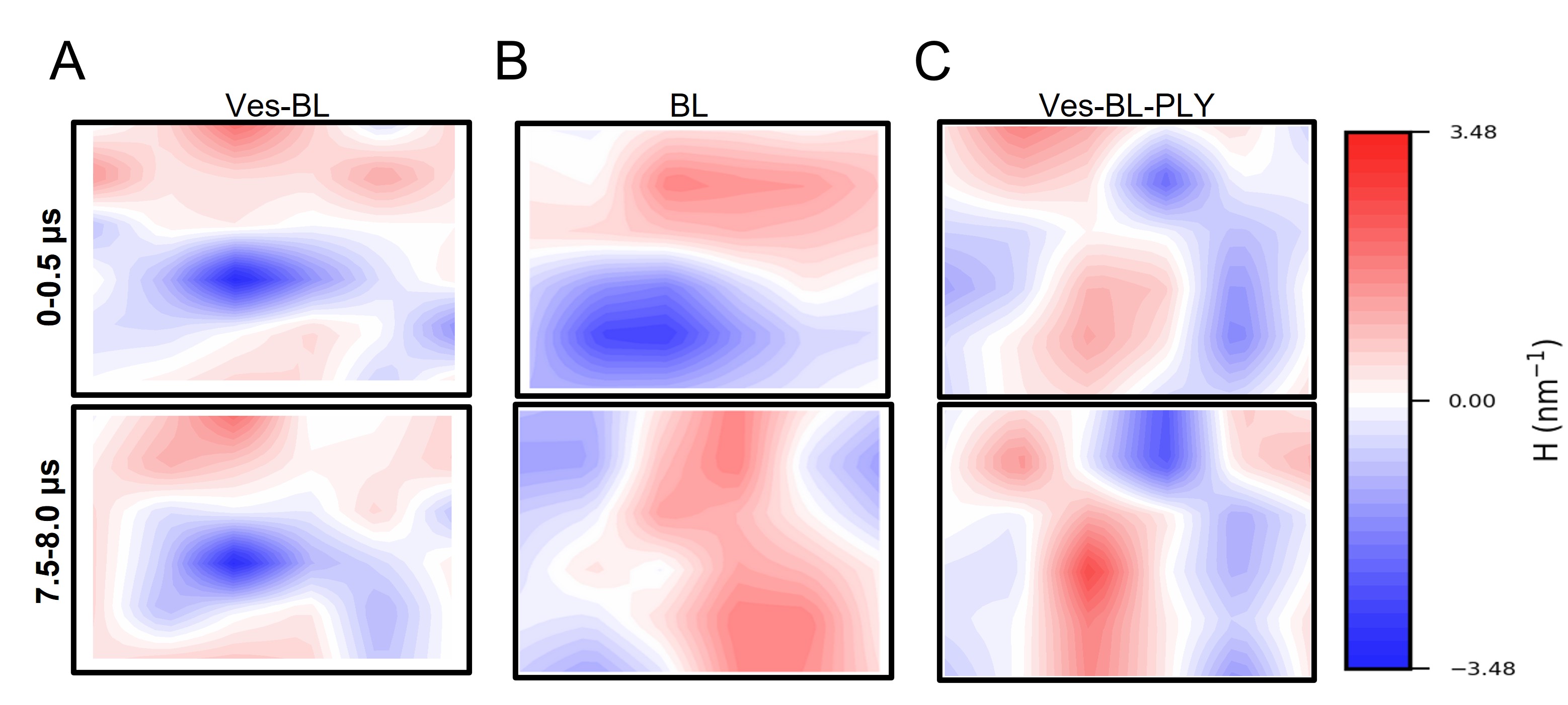

### Figure S3

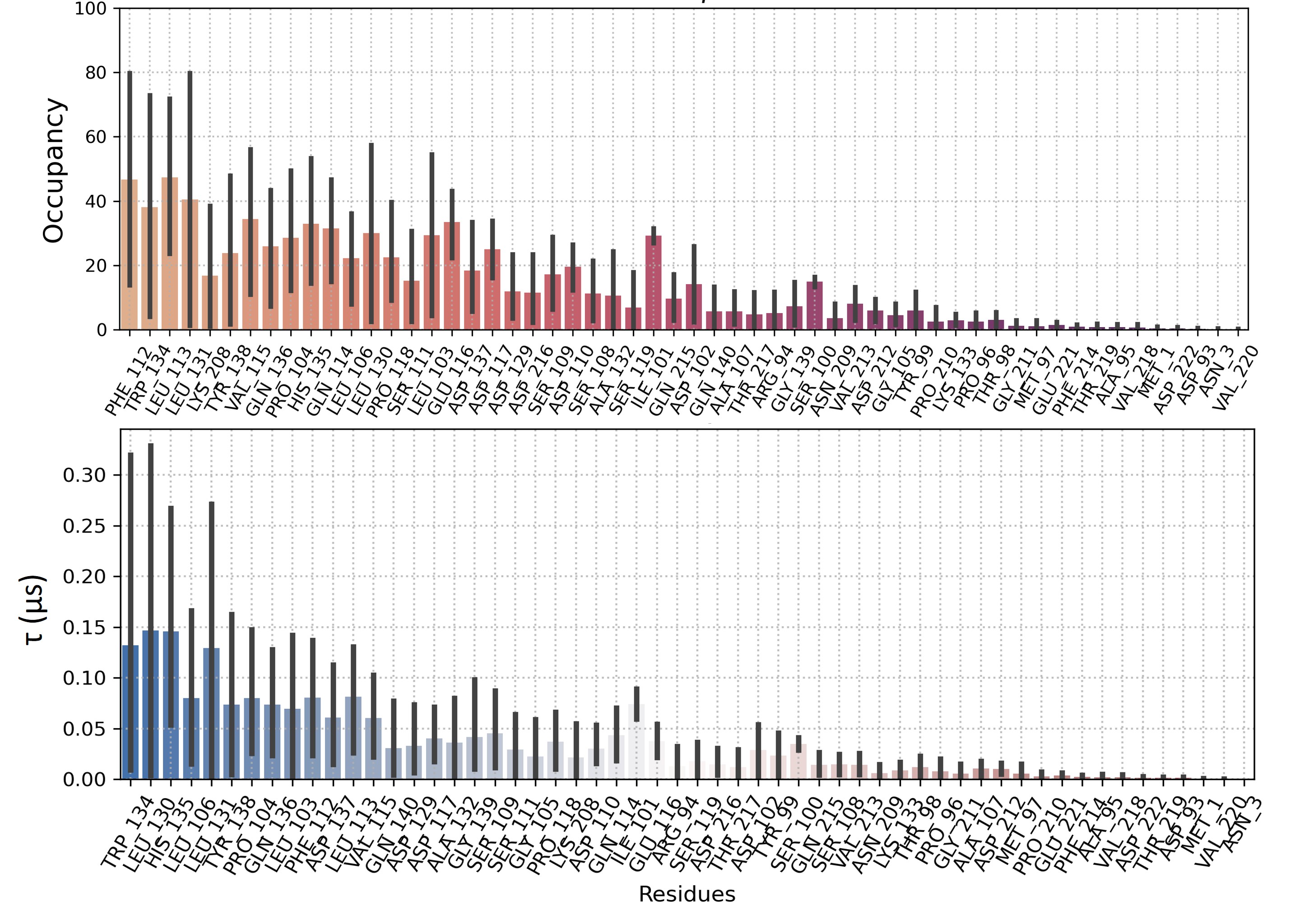

### Figure S4

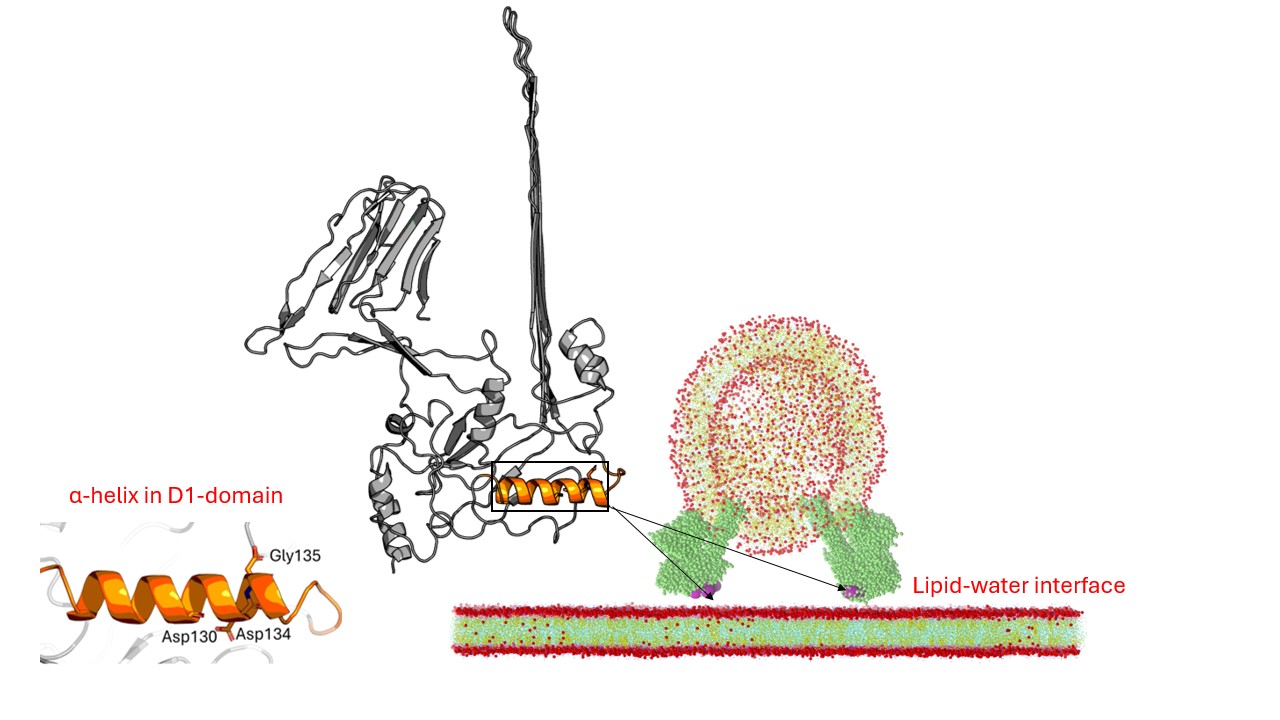

### Figure S5

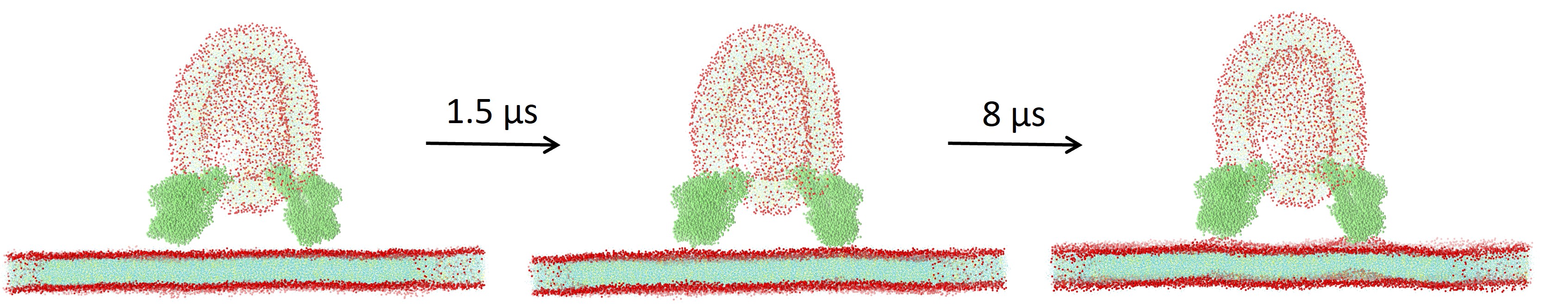

### Figure S6

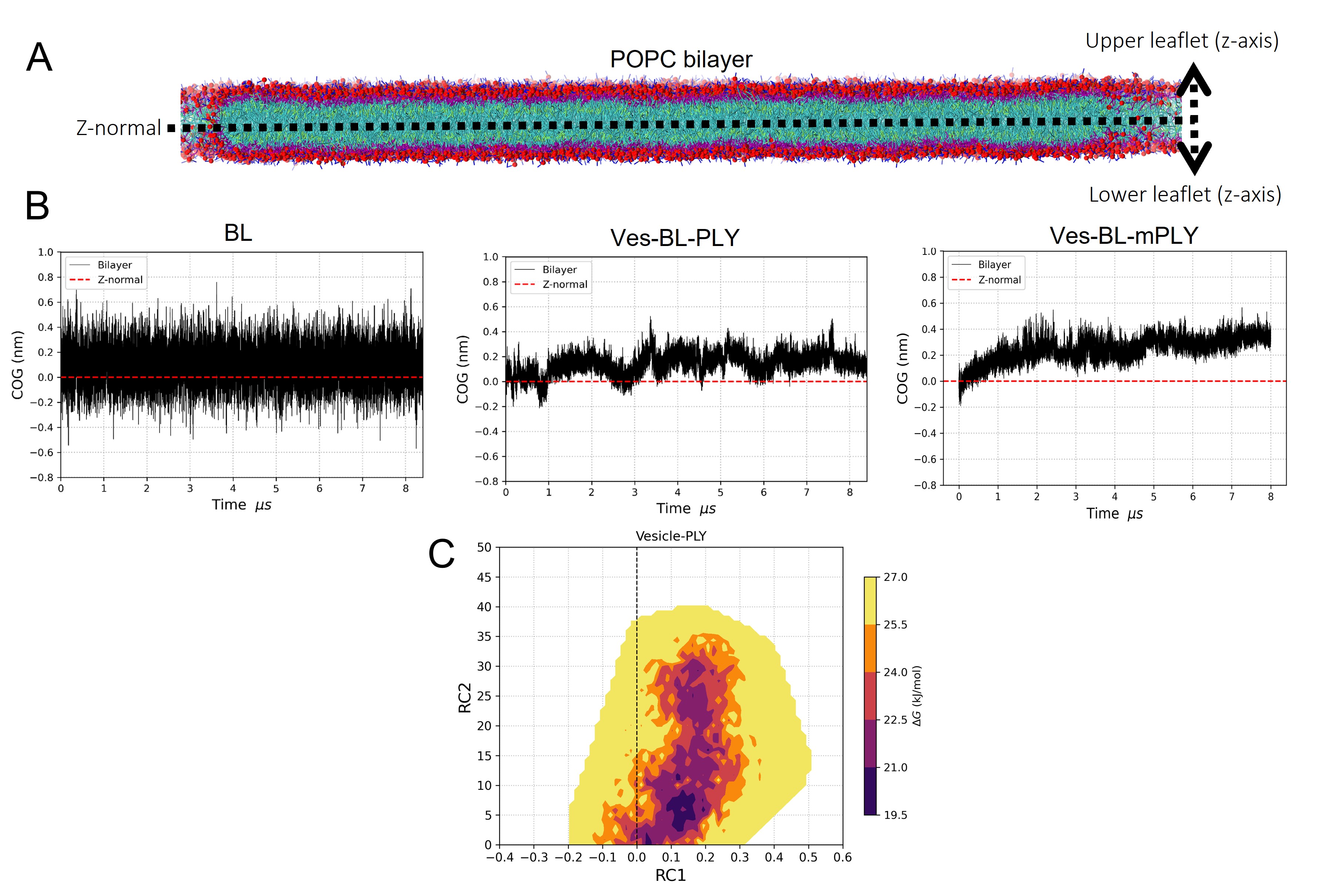

### Figure S7

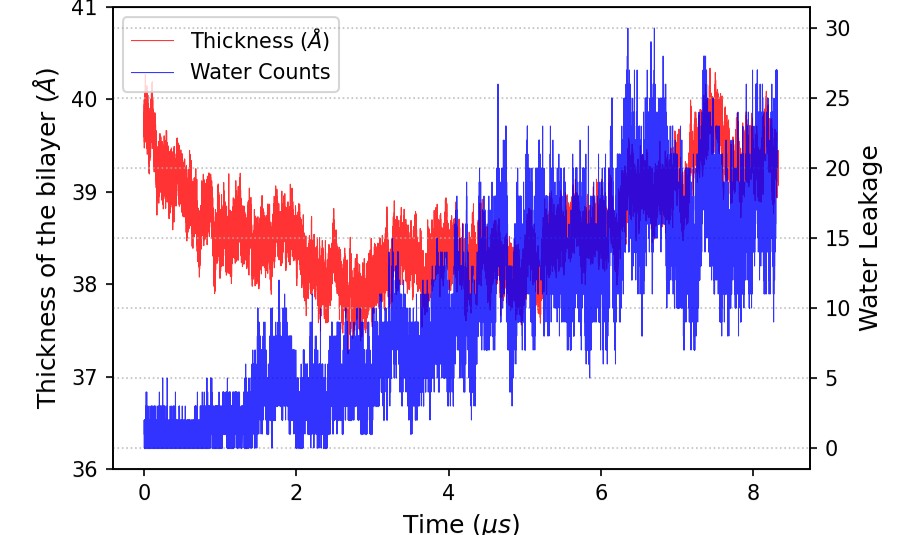

### Figure S8

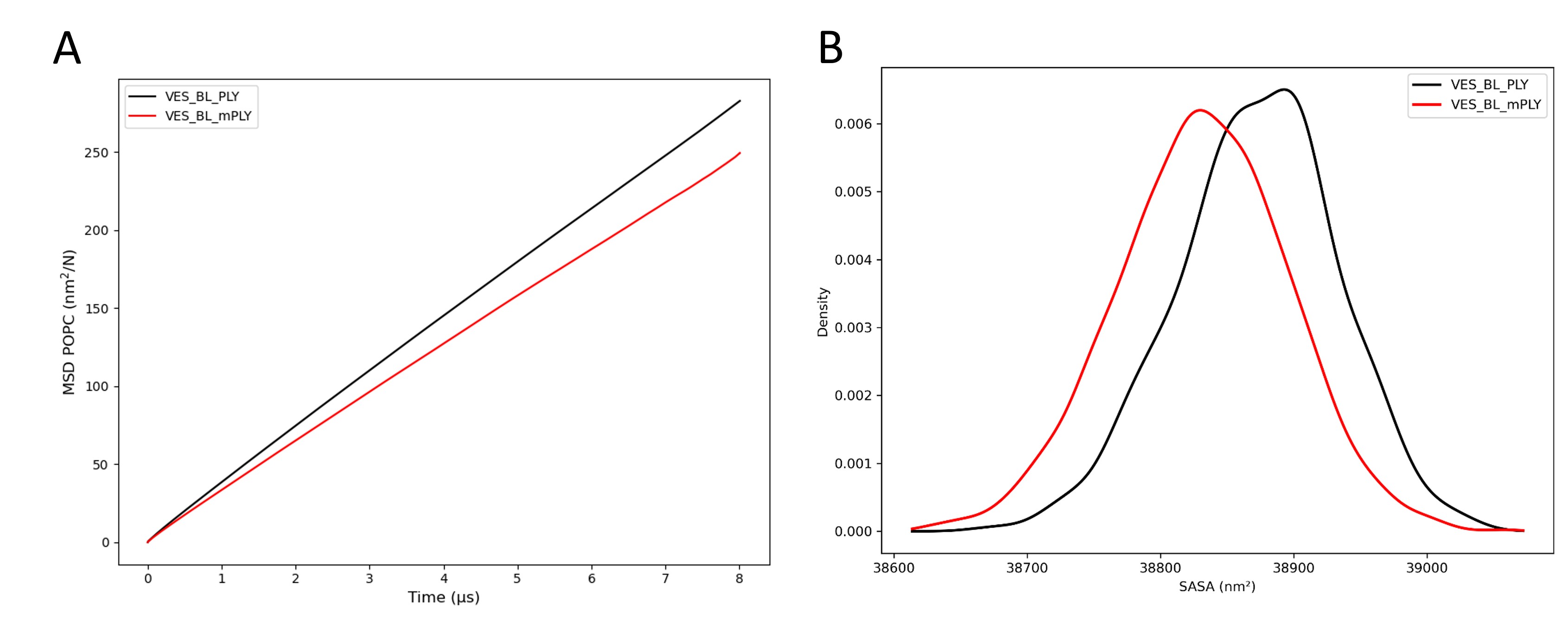
